# The plant calcium-dependent protein kinase CPK3 phosphorylates REM1.3 to restrict viral infection

**DOI:** 10.1101/205765

**Authors:** Artemis Perraki, Julien Gronnier, Paul Gouguet, Marie Boudsocq, Anne-Flore Deroubaix, Vincent Simon, Sylvie German-Retana, Cyril Zipfel, Emmanuelle Bayer, Sébastien Mongrand, Véronique Germain

## Abstract

Plants respond to pathogens through dynamic regulation of plasma membrane-bound signaling pathways. To date, how the plant plasma membrane is involved in responses to viruses is mostly unknown. Here, we show that plant cells sense the Potato virus X (PVX) COAT PROTEIN and TRIPLE GENE BLOCK 1 proteins and subsequently trigger the activation of a membrane-bound calcium-dependent kinase. We show that the *Arabidopsis thaliana* CALCIUM-DEPENDENT PROTEIN KINASE 3-interacts with group 1 REMORINs *in vivo*, phosphorylates the intrinsically disordered N-terminal domain of the Group 1 REMORIN REM1.3, and restricts PVX cell-to-cell movement. REM1.3-s phospho-status defines its plasma membrane nanodomain organization and is crucial for REM1.3-dependent restriction of PVX cell-to-cell movement by regulation of callose deposition at plasmodesmata. This study unveils plasma membrane nanodomain-associated molecular events underlying the plant immune response to viruses.

## Introduction

The cell plasma membrane (PM) constitutes a regulatory hub for information processing where intrinsic and extrinsic signals are perceived, integrated and relayed ^1^. PM proteins and lipids dynamically associate with each other, creating specialized sub-compartment that regulate the cellular responses in space and time ^2–4^. In animal, modeling of a PM-bound receptor and its downstream interactor before and after ligand perception, suggests that PM-partitioning into nanodomains improves the reliability of cell signaling ^5^. In plants, the immune and growth receptors FLS2 and BRI1 are divided into context-specific nanodomains, allowing them to regulate distinct signaling pools at PM, despite sharing several signaling components ^6^. The REMORIN (REM) family is probably one of the best-characterized PM nanodomain-associated proteins in plants ^6–11^. The association of REMs to the PM is mediated by a short sequence at the extremity of the C-terminus of the protein, called REM-CA (REMORIN C-terminal Anchor) ^12 13^. The REM C-terminal domain, contain a predicted coiled-coil (residues 117-152) which is thought to regulate REM oligomerization ^10, 13, 14^ and may be involved in regulating REM spatial organization at the PM. Members of the REM family have been associated with plant responses to biotic ^8, 15, 16^, abiotic stress ^17 18^ and developmental clues ^11^ and current view suggest they could regulate signaling events through nanodomain association. However, the molecular mechanisms leading to REM downstream events remain elusive. Several REM proteins have been identified as components of the plasmodesmata-plasma membrane subcompartment (PD-PM) ^7, 19, 20^. PD are membranous nanopores, crossing the plant cell wall and enabling cytoplasmic, endoplasmic reticulum and PM continuity between adjacent cells. They regulate the intercellular transport of proteins and small molecules during development and defense ^21 22^. The PD-PM is a particular subcompartment of the PM which displays a unique molecular composition and is enriched in sterols ^23^. The movement of macromolecules through PD can be tightly controlled through modulation of the PD size-exclusion limit (SEL) *via* hypo- or hyper-accumulation of callose at the PD neck region ^24–26^. Overexpression of GRAIN SETTING DEFECT 1 (GSD1), encoding a phylogenetic group 6 REM protein from rice, restricts PD aperture and transport of photoassimilates ^20^.

PD are also the only route available for plant viruses to spread from cell-to-cell. *Potato virus X* (PVX) promotes its cell-to-cell movement *via* modification of PD permeability ^27^ through the action of TRIPLE GENE BLOCK PROTEIN 1 (TGBp1) and callose removal ^28^. Overexpression of StREM1.3 (*Solanum tuberosum* REM from group 1b, homolog 3 ^29^), further referred as REM1.3, hampers TGBp1 ability to increase PD permeability ^30^. How is REM1.3 obstructing TGBp1 action is still unknown but its lateral organization into nanodomains at the PM is directly linked with REM1.3 ability to restrict PVX movement and regulate PD conductance ^31^.

REM1.3 was the first REM family member discovered and initially described as a protein phosphorylated upon treatment with oligogalacturonides, which are plant cell wall components and elicitors of plant defense ^32, 33^. The biological relevance of REM phosphorylation is not known, yet reports of different REM phospho-statuses suggest that the activity of these proteins could be regulated by phosphorylation during plant-microbe interactions ^14, 15, 34, 35^.

In the present paper, we show that phosphorylation of REM dictates its membrane dynamic and antiviral defense by the reduction of PD permeability. Our data points toward a model whereby viral proteins such as the Coat Protein (CP), TGBp1 from PVX and 30K proteins from *Tobacco mosaic virus* (TMV) elicit the activation of protein kinase(s), which in turn phosphorylate(s) REM1.3 at its N-terminal domain. In turn, REM1.3’s phospho-status regulates its spatial-temporal organization at the PM and association with PD. The latter is associated with PD closure by induction of PD callose deposition at PD and restriction of viral cell-to-cell movement. Last, we further provide evidence that the membrane bound *Arabidopsis thaliana* CALCIUM-DEPENDENT PROTEIN KINASE 3 (CPK3) interacts with the taxonomic group 1b REMs ^7^ *in vivo*, phosphorylates REM1.3 *in vitro* and restricts PVX propagation in a REM-dependent manner. Collectively, this study brings valuable information about the involvement of PM nanodomains dynamics during the establishment of membrane-bound signaling processes.

## Results

### PVX triggers changes in REM1.3 membrane dynamic behavior and REM1.3 association with plasmodesmata

Group 1 and group 6 REM have been described as PD-localized protein regulating PD size-exclusion limit ^7 20 30^. REM1.3 plays a role in restricting PVX passage through the PD channels ^7 30^, whereas in the same time PVX movement proteins promote opening of PD ^36^. To study the potential function of REM1.3 at PD in response to PVX infection, we surveyed simultaneously PD callose content and REM1.3 PD localization in healthy or PVX-infected *N. benthamiana* transgenic lines overexpressing *YFP-REM1.3* ^37^ (Fig EV1). Our analysis firstly showed a significant increase in callose deposition in PVX-infected cells (Fig 1A,B). This finding suggests the recognition of PVX-encoded elicitor and the mobilization of a plant defense response leading to an increase of callose accumulation at PD pit fields.

**Figure 1.**
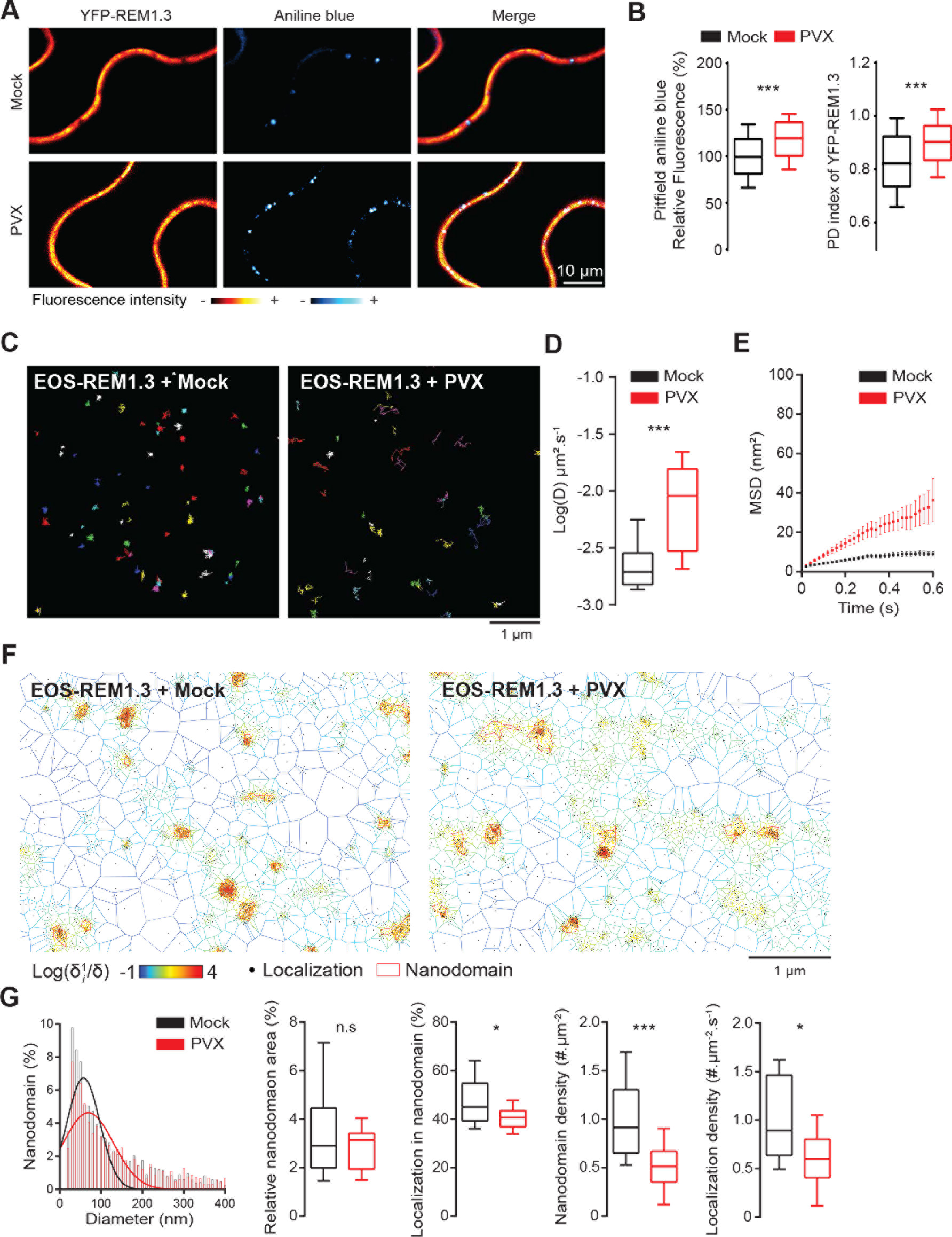
REM1.3 modulates Plasmodesmata callose accumulation and displays altered PM organization and dynamic following PVX infection. (A) Representative confocal images of aniline blue stained *N. benthamiana* leaf epidermal cells stably expressing YFP-REM1.3 in the absence (mock is infiltration with empty *A. tumefaciens*) or the presence of PVX at 2 days after infiltration (DAI). Color-coding indicates fluorescence intensity. (B) *left*, Pit field aniline blue fluorescence intensity was quantified by ImageJ as described in Fig EV1 and expressed as the percentage of the mock control. *right*, Quantification of the PD residency of YFP-REM1.3 in the absence (mock) and in the presence of PVX using the PD index ^25^ as described in Fig EV1. Graphs represent quantifications from 3 independent biological experiments. At least 15 cells per condition were analyzed per experiment. Significant differences were determined by Mann-Whitney comparisons test *** p<0.001. (C) Super-resolved trajectories of EOS-REM1.3 molecules in the PM plane in the absence (Mock) and presence of PVX obtained by high-resolution microscopy spt-PALM. (D) Diffusion coefficients (D) of EOS-REM1.3 expressed as log(D) in the absence (Mock) and presence of PVX. Statistical significances were assessed by Mann-Whitney test *** p<0.001 using data collected over two independents experiments. (E) Mean Square Displacement (MSD) over time for the global trajectories of EOS-REM1.3 followed during at least 600 ms reflecting two independent experiments. (F) Live PALM analysis of EOS-REM1.3 localization in the absence (mock) and presence of PVX by tessellation-based automatic segmentation of super-resolution images. (G) Computation of EOS-REM1.3 single molecule organization features based on tessellation-based automatic segmentation images. For REM1.3 nanodomain size distribution for the indicated conditions, the Gaussian fits in absence (mock) and presence of PVX are indicated by lines. Total nanodomain area is expressed as percentage of the total PM surface. Percentage of EOS-REM1.3 molecules localizing into nanodomains, relative to all molecules observed. Localization density refers to the number of molecules observed per μm^2^ per second. Statistics were performed on at least 10 data sets per condition, from two independent experiments. Significant differences were determined by Mann-Whitney test * p<0.05, *** p<0.001.

Since protein activation is often linked to changes in subcellular localization, even to specific PM sub-compartments ^2, 38^, we next examined whether PVX infection triggers changes in REM1.3 PD localization. Calculation of the PD index (ratio between fluorescence intensity of YFP-REM1.3 at the aniline-labeled PD pit field and fluorescence at the PM around the pit field ^25^, Fig EV1) surprisingly showed that despite its direct role on PD regulation YFP-REM1.3 is not enriched in the PD region of healthy *N. benthamiana* epidermal cells. Interestingly, we however reproducibly observed a slight increase of YFP-REM1.3 PD index upon PVX infection (Fig 1A,B) suggesting that PVX perception modulates REM1.3 localization and association with the PD pit fields.

To gain further insights into REM1.3 dynamic localization at the PM upon PVX infection, we applied single-particle tracking Photoactivated Localization Microscopy in Variable Angle Epifluorescence Microscopy mode (spt-PALM VAEM) in living *N. benthamiana* cells in absence or presence of PVX. By this approach, we recently studied the protein organization and mobility parameters of single EOS-REM1.3 molecules in non-infected conditions and found that EOS-REM1.3 displays a mostly immobile and confined PM localization pattern, as commonly observed for plant membrane-associated proteins (Fig 1C-E) ^39 31^. Reminiscent of these data, previous studies using different techniques described REM-associated PM domains to be predominantly stable to lateral diffusion ^31, 39, 40^. Analysis of PVX-infected cells demonstrated an increase of EOS-REM1.3 diffusion coefficient(D) and mean square displacement (MSD), reflecting an increase of REM1.3 mobility (Fig 1C-E). We next compared the supra-molecular organization of EOS-REM1.3 by Voronoï tessellation of live PALM data ^31 41^ in mock and PVX infected conditions (Fig 1F). Computation of EOS-REM1.3 single molecule organization features demonstrated a modulation of REM1.3 nanodomain-organization upon PVX infection (Fig 1G). Following PVX infection, the EOS-REM1.3-formed nanodomains are bigger in size, and there is a slight decrease of the proportion of molecules that localizes into nanodomains as well as a significant decrease in the number of nanodomains formed. Overall, in both conditions, EOS-REM1.3 nanodomains represented similar proportions of the total PM surface. Additionally, a significant decrease in the localization density (number of molecules observed per μm^2^ per s), shows that upon PVX infection, there is less protein at PM level, which could reflect relocalization of molecules to other compartments. Overall, the changes of REM1.3 distribution at the PM under PVX infection *i.e.* enrichment of YFP-REM1.3 in the pit field regions, the increase of REM1.3’s mobility and the modulation of REM1.3 nanodomain organization, suggest that the plant cell might modulate PD-PM nanodomain dynamics to circumvent PVX infection.

### Perception of PVX proteins by plant cells leads to the activation of kinase(s) phosphorylating REM1.3

*REM1.3* overexpression was shown to restrict PVX spreading in both *Solanum lycopersicum ^7^* and *Nicotiana benthamiana* ^30, 31^. As REM1.3 was originally discovered as a PM-associated phosphorylated protein ^33^, we asked whether we could detect kinase activity in *N. benthamiana* extracts leading to REM1.3 phosphorylation and whether this activity was induced by PVX presence in the cells. Equal protein amounts of microsomal and soluble extracts from *N. benthamiana* leaves were used as a kinase source to phosphorylate affinity-purified full-length 6His-REM1.3 in an *in vitro* kinase assay in the presence of ATP [γ-^33^P]. Autoradiography revealed the presence of a clear band corresponding to a phosphorylated form of 6His-REM1.3 by kinase(s) present in the microsomal fraction (Fig 2A). The intensity of this band was reduced and completely abolished by competition with cold ATP, but not cold AMP, indicating that this was a valid experimental set-up to study a genuine transphosphorylation event (Fig EV2A). Phosphorylation of 6His-REM1.3 was almost not detectable in soluble fractions, representing cytosolic kinases (Fig 2A).

**Figure 2.**
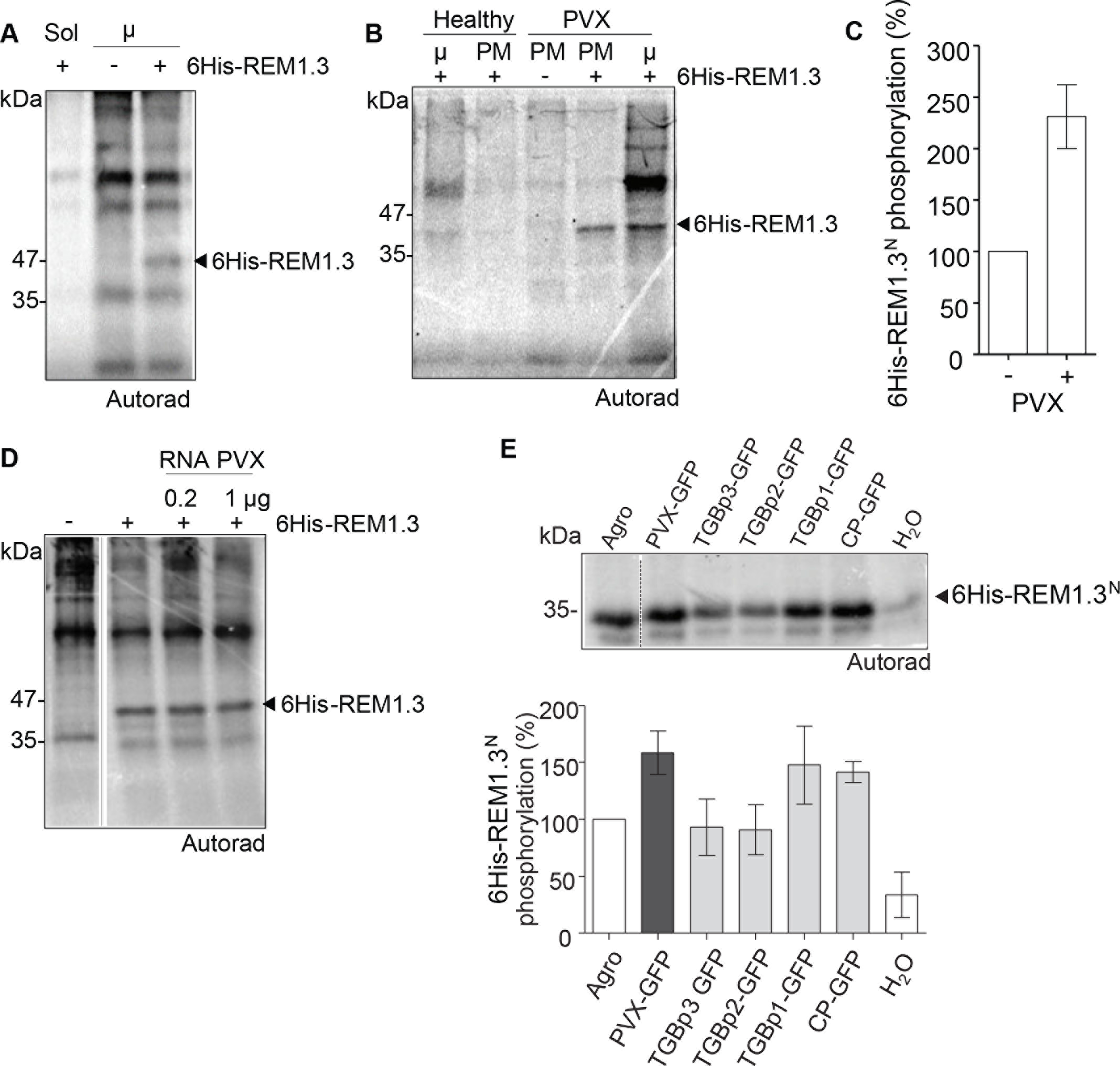
PVX and viral proteins induce REM1.3 phosphorylation in its N-terminal domain. (A, B) In *vitro* protein phosphorylation assays were performed by incubation of recombinant affinity-purified 6His-REM1.3 and *N. benthamiana* extracts with [γ-^33^P]-ATP. The samples were run on SDS-PAGE gels and developed by autoradiography. Soluble (Sol) or microsomal (μ) extracts of healthy leaves in (A), or microsomal and PM extracts from healthy and PVX-infected plants in (B) were used. (C) *In vitro* phosphorylation of 6His-REM1.3^N^ by leaf microsomal extracts of healthy or PVX-infected *N.benthamiana* leaves. Bars show the quantification of phosphorylated 6His-REM1.3^N^ bands from 5 independent repeats. (D) *In vitro* phosphorylation of 6His-REM1.3 by leaf microsomal extracts in the presence of total RNA extracts from PVX-infected leaves. (E) 6His-REM1.3^N^ phosphorylation by microsomal extracts infected with PVX-GFP or expressing the indicated viral proteins at 4 DAI. Leaves expressing GFP alone, infiltrated with water or with *A. tumefaciens* strain GV3101 alone served as controls. Graph represents the quantification of 6His-REM1.3^N^ bands from 3 independent repeats, as a percentage of the activity induced by PVX-GFP. In all experiments 10μg of total protein extracts and 1μg of affinity purified 6His-REM1.3 or 6His-REM1.3^N^ were loaded per lane.

*In silico* analysis predicted phosphorylation sites throughout REM1.3 sequence (Diphos, DEPP and NETPHOS prediction software). In agreement with the location of the sites presenting the highest phosphorylation potential, we experimentally found that REM1.3 was phosphorylated in its N-terminal domain (residues 1–116, hereby 6His:REM1.3^N^) whereas the C-terminal domain (residues 117–198, hereby 6His:REM1.3^C^) did not present any detectable phosphorylation (Fig EV2B,C). We next tested whether PVX activates the membrane-associated kinase(s) that phosphorylate(s) REM1.3. Our results unveiled that microsomal and PM fractions extracted from symptomatic PVX-infected leaves promoted higher levels of 6His-REM1.3 phosphorylation compared to extracts from non-infected plants (Fig 2B,C).

Studies have shown that functionally different viral components, such as virus-encoded proteins and double stranded RNA, can trigger plant defense responses ^42 43 44, 45 46, 47^. We examined whether the PVX genome in its free form was an eliciting signal for kinase activation and found that the addition of total RNAs extracted from PVX-infected plants in the kinase reaction mix did not alter the levels of 6His-REM1.3 phosphorylation (Fig 2D). We then examined whether the sole expression of individual viral movement proteins was sufficient to trigger REM1.3 phosphorylation. Importantly, our results demonstrated that the expression of TGBp1 and Coat Protein (CP) fused to GFP triggered the strongest levels of 6His-REM1.3^N^ phosphorylation to the same extent as the full PVX-GFP construct (Fig 2E). TGBp2 and TGBp3 proteins also induced 6His-REM1.3^N^ phosphorylation but to a lesser extent than TGBp1, CP, or full PVX-GFP. Quantification of three independent biological experiments shows that TGBp2 and TGBp3-induced 6His-REM1.3N phosphorylation at levels comparable to the control mock experiments (Fig 2E). Expression of a TGBp1-deleted version of PVX (PVXΔTGBp1) decreased 6His-REM1.3 phosphorylation levels compared to wild-type PVX extracts, but still to a greater level than GFP alone (Fig EV2D). Infiltration of the empty *Agrobacterium* strain alone induced stronger phosphorylation on 6His-REM1.3^N^ in *N. benthamiana* epidermal cells than the water control (Fig 2E) supporting that REM can be differentially phosphorylated during plant-microbe interactions^8^. Interestingly, expression of the 30K-RFP protein from *Tobacco mosaic virus* (TMV) also induced REM phosphorylation (Fig EV2E), suggesting a broader role of REM-mediated plant response to different genera of viruses. In good agreement with this, REM1.3 was shown to interfere with the ability of both PVX-TGBp1 and TMV-30K to increase PD permeability ^30^. Concomitantly, overexpression of REM1.3 was found to restrict TMV-GFP cell-to-cell movement in *N. benthamiana* epidermal cells (Fig EV3A).

Collectively these results suggest that the triggering of REM1.3 phosphorylation by the perception of viral proteins by plant cells might regulate REM1.3 function in PD permeability regulation.

### Phosphorylation of REM1.3 on critical residues of its N-terminal domain regulates its function in restricting PVX spreading via PD aperture modulation

Since differential phosphorylation of REM occurs upon PVX infection, we next aimed to functionally characterize the importance of REM1.3 phosphorylation for the regulation of PVX cell-to-cell movement. Despite our efforts, the identification of *in vivo* phosphorylation sites of REM1.3 appeared technically challenging and remained unsuccessful to this day. *In silico* predictions and *in vitro* kinase assays however showed that REM1.3^N^ displays regions of intrinsic disorder and presents the highest potential of phosphorylation (Fig 2 C,E, and Fig 3A). For functional characterization, we selected the three putative phosphorylation Ser/Thr sites present in REM1.3^N^, namely S74, T86 and S91, that presented high scores of phosphorylation prediction in intrinsic disorder regions (Fig 3A). S74 and S91 are conserved across the phylogenetic group 1b of REM proteins, suggesting functional redundancy (Fig EV5A) ^29, 48^. S74 and S91 were the analogous residues identified as phosphorylated *in vivo* in the group 1b REM AtREM1.3 (At2g45820) of *Arabidopsis thaliana* (hereby Arabidopsis) in a stimuli-dependent manner ^34, 35, 48^. Biochemical analysis showed that alpha-1,4-poly-D-galacturonic acid (PGA)-induced phosphorylation of StREM1.3 occurs on T32, S74 and T86 ^49^. T86 is not conserved in Arabidopsis but it is conserved in Solanaceae REM proteins, such as in *N. benthamiana* (Fig EV5A). By an *in vitro* kinase assay, we were able to demonstrate that S74, T86 and S91 are true phosphorylated residues of REM1.3, since substitution of S74, T86 and S91 to the non-phosphorylable Aspartic acid (D), generating the 6His-REM1.3^DDD^ mutant abolished REM phosphorylation by the PVX-activated kinase(s) (Fig 3B,C).

To functionally characterize the relevance of different REM1.3 phospho-statuses in the context of PVX-GFP propagation and PD-aperture regulation, we generated RFP- and YFP-fused REM1.3 mutants substituted at those sites either with Aspartic acid (REM1.3^DDD^) to mimic constitutive phosphorylation (hereby termed phosphomimetic mutant) or with Alanine (REM1.3^AAA^) representing null-phosphorylated state (hereby termed phosphodead mutant). Infection assays in *N. benthamiana* demonstrated that the phosphodead mutant lost ability to induce the restriction of PVX-GFP spreading, while the phosphomimetic mutant maintained this ability (Fig 3D). In good agreement with the TMV-30K-stimulated phosphorylation of REM1.3 presented in Fig EV2E, we confirmed that TMV-GFP propagation was also affected by the phospho-status of REM1.3 to the same extent as for PVX-GFP (Fig EV3A).

**Figure 3.**
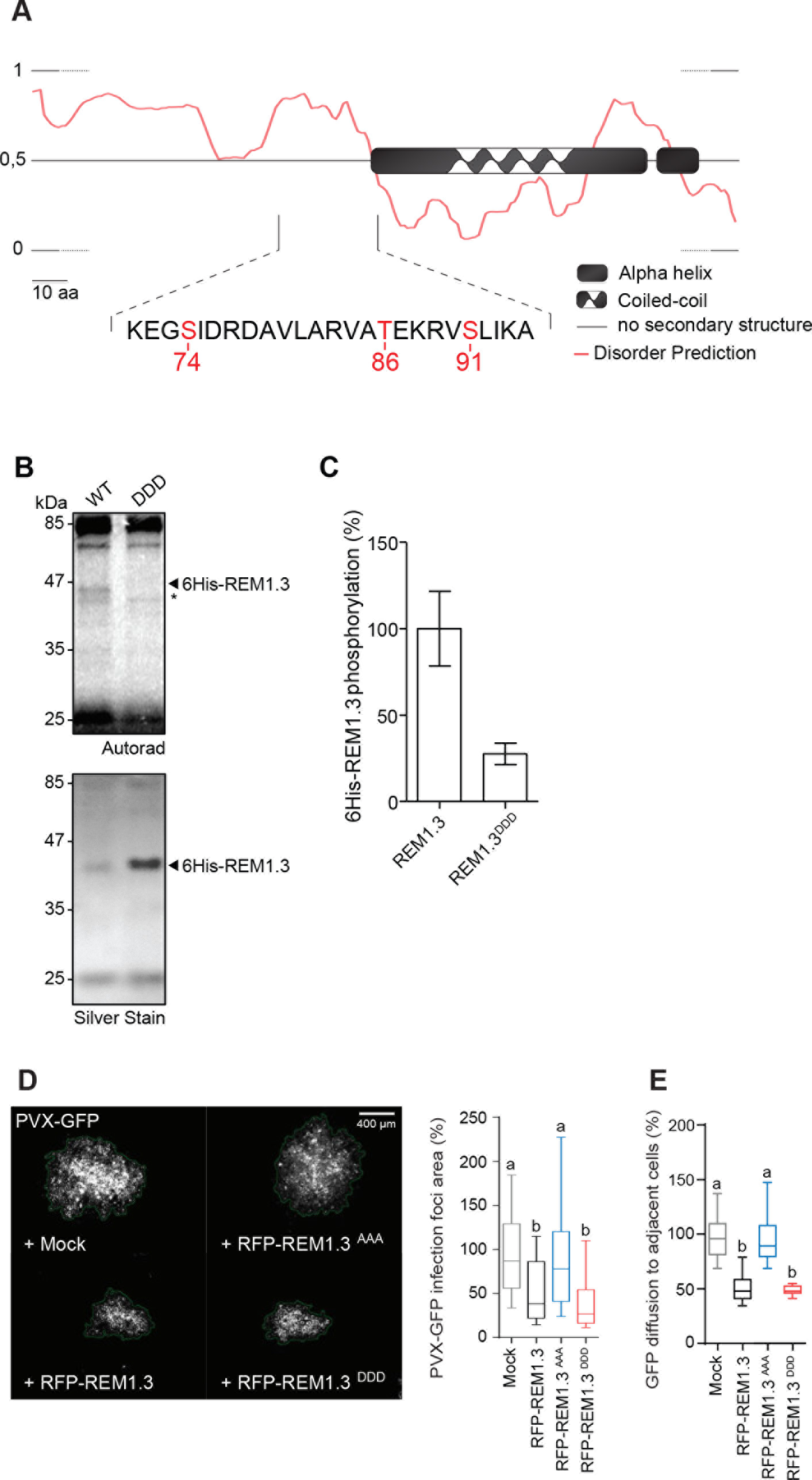
Mutational analysis reveals three critical phospho-residues required for REM1.3 regulation of PVX-GFP propagation and PD conductance. (A) *In silico* analysis of REM1.3 protein sequence. Prediction of putative phosphorylation sites was performed by Diphos, DEPP and NETPHOS coupled with published MS data. Predictions highlight three residues S74, T86 and S91 with high probability to be phosphorylated. Disordered prediction was performed by pDONR VL XT. Numbers indicate amino acid position.(B) *In vitro* kinase assay on recombinant affinity purified 6His-REM1.3 or 6His-REM1.3DDD by incubation with [γ-^33^P]-ATP and microsomal extracts of PVX-infected *N. benthamiana* leaves, as described in Figure 2. In (C), graph represents the relative quantifications from 4 independent reactions, using WT signal as a reference. Asterisk * indicates phosphorylation of a *N. benthamiana* protein of close molecular weight not detected by silver staining. (D) Representative epifluorescence microscopy images of PVX-GFP infection foci on *N. benthamiana* leaf epidermal cells at 5 DAI. Graph represents the mean relative foci area of REM1.3 or phosphovariants compared to mock control (co-infiltration of PVX-GFP with an empty *A. tumefaciens* strain). Approximately 250 foci per condition from 5 independent biological repeats were measured Letters indicate significant differences revealed by Dunn’s multiple comparisons test p<0.001. (E) GFP diffusion to neighbor cells was estimated by epifluorescence microscopy at 5 DAI in *N. benthamiana* cells expressing REM1.3 or phosphomutants. Measurements from 3 independent biological repeats were normalized to mock control (co-infiltration with an empty *A. tumefaciens* strain). Letters indicate significant differences determined by Dunn’s multiple comparisons test p<0.001.

We then analyzed the capacity of REM1.3 phosphomutants to regulate PD aperture in the absence of viral infection. As previously described ^30, 31^, RFP-REM1.3 reduces the PD size-exclusion limit as measured by free GFP diffusion to neighboring cells (Fig 3E). Detailed phenotypic analysis of REM1.3 phosphorylation mutants demonstrated that the phosphomimetic mutant recapitulated REM1.3 activity towards PD-aperture regulation, while the phosphodead mutant did not (Fig 3E). Altogether, these results provide strong evidence that REM1.3’s phosphorylation state at the evolutionarily conserved positions of S74, T86 and S91 is linked to its function in controlling viral infection and PD conductance.

### REM1.3 phospho-status modulates its dynamic lateral segregation in PM and PD sub-compartments

Upon PVX infection we observed a modulation of REM1.3 PD-association and PM dynamics (Fig 1), linked to REM1.3 phosphorylation (Fig 2) that is required for REM1.3 function against PVX infection (Fig 3). Hence, we asked whether different REM1.3 phospho-statuses might regulate its lateral organization at the PM and PD compartments in the absence of PVX. We examined the enrichment of REM1.3 phosphomutants at the PD pit fields, previously calculated by the PD index (Fig 1A,B) and found that similarly to YFP-REM1.3, none of the phosphomutants appeared enriched at the pit field level (Fig 4A, C). The phosphodead mutant appeared statistically more excluded than YFP-REM1.3, whereas the phosphomimic mutant displayed an increase of its PD index (Fig 4C), reminiscent of the REM1.3 localization phenotype under PVX infection (Fig 1A,B). Importantly, REM1.3 phosphomutants’s association with PD was directly correlated with callose content at PD (Fig 4B). These observations reinforced the hypothesis that REM1.3-mediated increase of callose levels at the PD is associated with a dynamic and phosphorylation dependent redistribution of REM1.3 to the PD surroundings.

**Figure 4.**
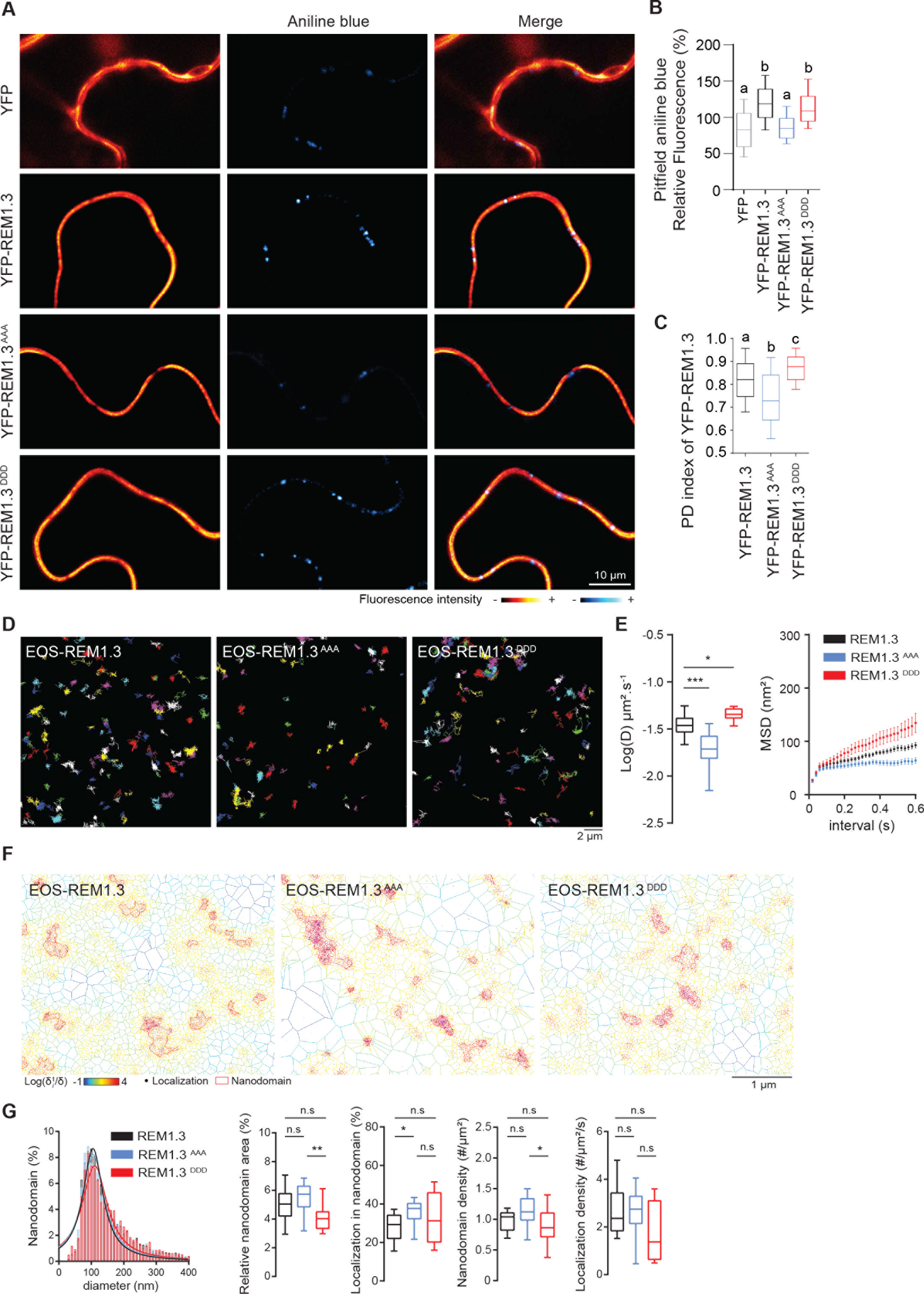
REM1.3 dynamic localization in PD and PM nanodomains is regulated by its phospho-status. (A) Representative confocal mages showing aniline blue staining of callose deposition at the PD pitfields in *N. benthamiana* leaf epidermal cells expressing YFP-REM1.3 or phosphovariants. Color-coding indicates fluorescence intensity. (B) Graphs show aniline blue fluorescence intensities in cells transiently expressing YFP-REM1.3 and phosphomutants relative to control cells expressing YFP alone. Three independent biological experiments were performed and at least 15 cells per condition and per experiment were analyzed. Letter indicate significant differences revealed by Dunn’s multiple comparisons test p,0.001. (C) PD index of YFP-REM1.3 phosphomutants was calculated as described in Figure EV1. Graphs present quantifications from 3 independent biological experiments. Letter indicate significant differences revealed by Dunn’s multiple comparisons test p,0.002. (D) Super-resolved trajectories of EOS-REM1.3, and phosphomutants, transiently expressed in *N. benthamiana* cells, observed by spt-PALM. Scale bars, 2 μm. (E) Distribution of diffusion coefficients (D) represented as log(D) of the different fusion proteins. Mean Square Displacement (MSD) over time for the global trajectories of each EOS-REM1.3 construct followed during at least 600ms. 27 cells for EOS-REM1.3, 15 cells for EOS-REM1.3^AAA^ and 17 cells for EOS-REM1.3^DDD^ were analyzed in 3 independent experiments. Statistical analysis was performed by Mann-Whitney test * p<0.05 ** p<0.01. (F) Live PALM analysis of EOS-REM1.3 phosphomutants by tessellation-based automatic segmentation of super-resolution images. (G) Computation of EOS-REM1.3 and phosphomutants single molecule organization features based on tessellation-based automatic segmentation images. For REM1.3 and phosphomutants nanodomain size distribution and the Gaussian fits are indicated. Total nanodomain area is expressed as percentage of the total PM surface. Percentage of EOS-REM1.3 molecules localizing into nanodomains, relative to all molecules observed. Localization density refers to the number of molecules observed per μm^2^ per second. Statistics were performed on at least 13 data sets per condition extracted from 3 independent experiments. Statistical differences determined by Mann-Whitney test * p<0.05, ** p<0.01.

We examined by confocal microscopy the localization pattern of the phosphomutants fused to YFP. Both phosphodead (YFP-REM1.3^AAA^) and phosphomimetic (YFP-REM1.3^DDD^) mutants located exclusively at the PM, analogous to REM1.3 localization (Fig EV3B). We next used spt-PALM VAEM to characterize the localization and mobility behavior of the EOS-REM1.3 phosphomutants. The analysis of reconstructed trajectories and corresponding super-resolved localization maps indicated slight modifications of lateral mobility behavior between the phosphomutants (Fig 4 D, E). Quantification of the diffusion coefficient values (D) extracted for each individual molecule revealed that EOS-REM1.3^AAA^ displayed a more immobile behavior than EOS-REM1.3^DDD^ and EOS-REM1.3. Consistently, EOS-REM1.3^DDD^ exhibited a higher mobility illustrated by higher diffusion coefficient and mean square displacement values (Fig 4 D, E). Analysis of the supra-molecular organization of the phosphomutants by Voronoï tessellation (Fig 4F) firstly showed that all mutants displayed similar nanodomain size and localization density compared to EOS-REM1.3^WT^. Compared to EOS-REM1.3AAA, the EOS-REM1.3^DDD^ nanodomains occupied a smaller area of the total PM and their density in the PM plane appeared slightly reduced (Fig 4F,G). A higher number of nanodomains were formed with the EOS-REM1.3^AAA^ mutant. Hence, the phosphomimetic mutations favor a less confined and a more dynamic localization pattern of REM1.3 at the PM, reminiscent to the phenotype of EOS-REM1.3^WT^ in the context of PVX infection (Fig 1C,D). These results suggest that differential REM1.3 phosphorylation is involved in regulating REM1.3 mobility and PM domain organization and support the hypothesis that REM1.3 phosphorylation on S74, T86 and S91 reflects an ‘active form’ of the protein necessary for REM1.3-mediated defense signaling.

### AtCPK3 phosphorylates REM1.3

To gain more insights into the signaling processes leading to REM1.3 phosphorylation, we aimed to biochemically characterize the kinase(s) involved in the phosphorylation of REM1.3. Previous evidence suggested that the kinase(s) phosphorylating REM1.3 are membrane-associated (Fig 2) ^33^. We therefore biochemically analyzed the localization of the kinase(s) phosphorylating REM1.3. Plant material from healthy and PVX-GFP-infected leaves was cell-fractionated to obtain crude extracts, soluble and microsomal fractions ^50^ to perform *in vitro* kinase assays on REM1.3^N^. Analysis confirmed a maximal kinase activity in purified microsomes (Fig 5A, 2A). Since a kinase in close proximity with its substrate would enhance reaction kinetics ^51^ and signal fidelity ^52^, and given that REM1.3 is enriched in detergent-resistant membranes (DRM) ^7^, we investigated whether the kinase activity towards 6His-REM1.3 is enriched in this biochemical fraction. We included “control PM” (C-PM) preparations, submitted to discontinuous sucrose gradients but in the absence of Triton-X100 treatments ^53^. *In vitro* kinase assays on 6His-REM1.3^N^ showed that the kinase activity in C-PM was 5 times inferior than in freshly purified PM not submitted to the sucrose gradient, suggesting that the kinase is not stable during the overnight purification procedure. Only half of the specific activity of the kinase was found in DRMs compared to the C-PM fraction, indicating that the kinase(s) phosphorylating REM1.3 is (are) only partially located in the DRM fraction (Fig 5B).

**Figure 5.**
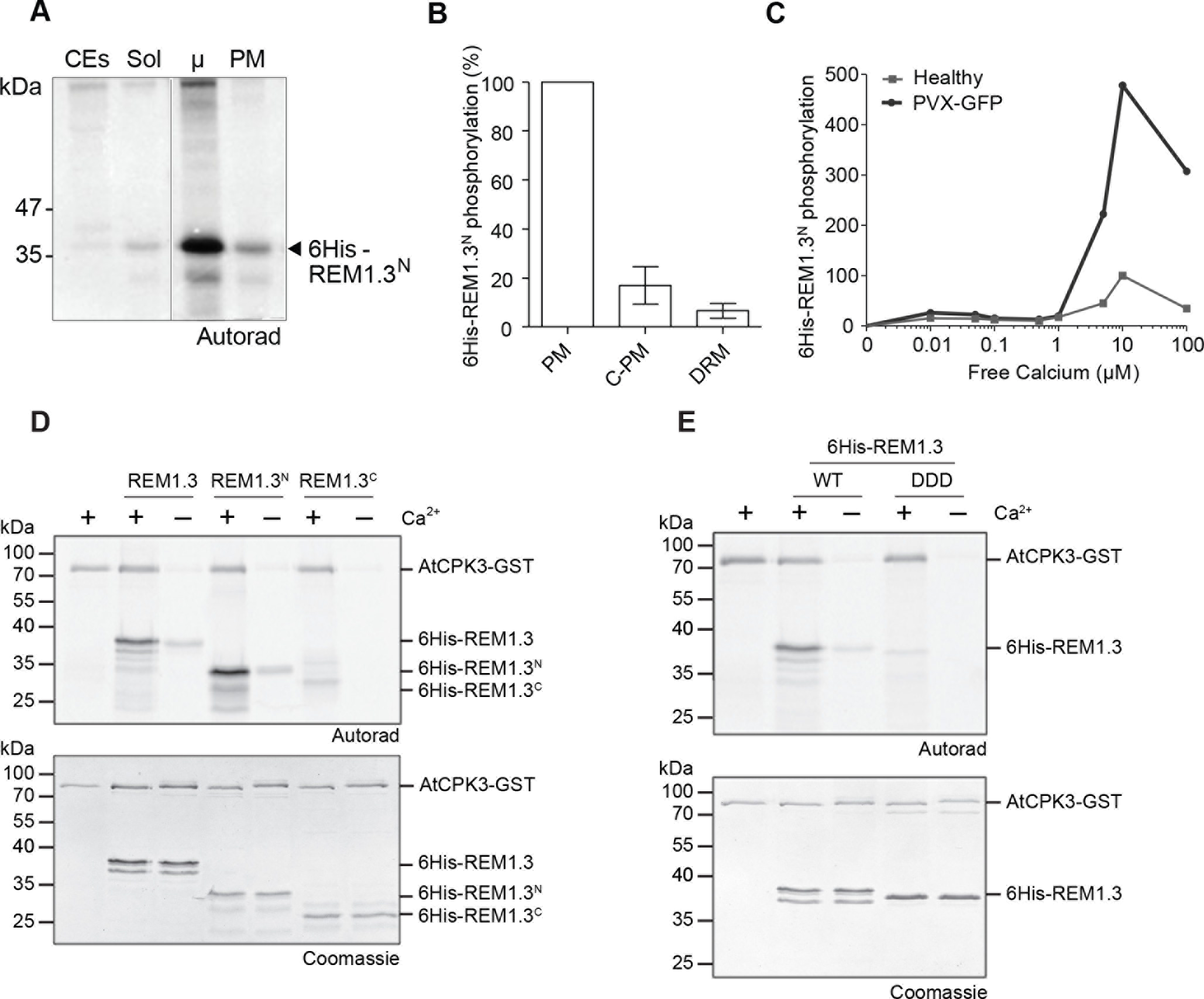
AtCPK3 phosphorylates REM1.3 in a calcium-dependent manner. (A, B) *In vitro* phosphorylation of purified 6His:REM1.3^N^ by kinase(s) from different cellular fractions of *N. benthamiana* leaves, CEs, leaf crude extracts; Sol, Soluble fraction; μ, microsomal fraction; PM, Plasma Membrane; C-PM: “Control-PM” is PM fraction not treated by TX100, but submitted to sucrose gradient; DRM, Detergent resistant membranes ^53^. The graph represents the relative quantification of 3 independent experiments normalized to the activity in the PM fraction +/− SEM. (C) Quantification of the calcium dose response of kinase activity on 6His-REM1.3^N^ phosphorylation by *N. benthamiana* microsomal extracts from healthy and PVX infected leaves. (D, E) Autoradiography gels show *in vitro* phosphorylation of 6His-REM1.3, 6His-REM1.3^N^ and 6His-REM1.3^C^ or 6His:REM1.3^DDD^ by affinity purified GST-AtCPK3 in the presence or the absence of Ca^2+^. Bands corresponding to autophosphorylation of AtCPK3-GST and transphosphorylation of 6His-REM1.3 variants are indicated. Gels were stained by coomassie blue to visualize protein loading.

To gain more information concerning the biochemical characteristics of the kinase phosphorylating REM1.3, we analyzed its activity in the presence of known inhibitors. We firstly tested staurosporine, ^54, 55^ a general inhibitor that prevents ATP binding to kinases and found inhibition of REM1.3 phosphorylation starting at very low concentrations (30 nM) (Fig EV4A). We further tested the effect of poly-L-lysine, described to stimulate the activity of the CK2 kinases and inhibit several CDPK kinases ^56, 57^. No significant differences on REM1.3 phosphorylation levels were observed under increasing concentrations of poly-L-lysine (Fig EV4B). The addition of the wide range of Ser/Thr phosphatases inhibitor β-glycerophosphate (BGP ^58^) to the reaction mix did not alter the levels of phosphorylated 6His-REM1.3, indicating that the observed data was due to the activation of kinase activity by PVX rather than by inhibition of phosphatases (Fig EV4B). Competition assays in the presence of cold AMP and GTP showed that only cold ATP even at 2 mM caused 20-fold depletion in [γ-^33^P] incorporation, suggesting that ATP is the major phosphoryl-donor for the kinase (Fig EV4B). Addition in the reaction mix of 0,2 mM of EGTA, a chelator of Ca^2+^, strongly inhibited the kinase activity suggesting that the kinase(s) phosphorylating REM1.3 in healthy leaves is calcium sensitive (Fig EV4C). Calcium is a conserved second messenger in signal transduction during biotic and abiotic stress. In plants, kinases harboring different calcium sensitivities can perceive calcium variations and translate them into downstream signaling activation ^59 60^. To determine whether the PVX-activated kinase phosphorylating REM1.3 is sensitive to calcium regulation, *in vitro* kinase assays from microsomes of healthy and PVX-infected *N. benthamiana* leaves were assayed in the presence of free calcium (Ca^2+^) concentrations ranging from 10 nM to 0,1 mM. Figure 5C shows that the kinase(s) displays a high sensitivity to calcium with an optimal activity in the presence of 10 μM of free Ca^2+^. At this concentration, a 5-fold increase of 6His-REM1.3^N^ phosphorylation was observed in PVX-infected leaves (Fig 5C). These experiments allowed us to narrow-down the kinase(s) phosphorylating REM1.3 after PVX infection to the group of membrane-bound Ca^2+^-dependent protein kinases ^59^. Plants possess three main families of calcium-regulated kinases: calmodulin-binding kinases (CBKs), calcineurin B-like-interacting protein kinases (CIPKs) and calcium-dependent protein kinases (CDPKs) ^58^. CDPKs have the unique feature of calcium sensing and responding activities in one single polypeptide, best characterized in the model plant *Arabidopsis thaliana ^59^*. Based on the measured calcium dose response (Fig 5C), we correlated the kinase phosphorylating REM1.3 in *N. benthamiana* with homologs of *Arabidopsis thaliana* subgroup II AtCPKs ^61^, and we aimed to capitalize on the knowledge of Arabidopsis CPKs to test REM1.3 phosphorylation. Among the characterized members of subgroup II AtCPKs, we selected the *Arabidopsis thaliana* AtCPK3 as a good candidate to test its putative role in REM1.3 phosphorylation, since previous proteomics studies in *Arabidopsis thaliana* have identified both AtCPK3 and AtREM1.3 as being enriched in PM, PD and DRM fractions ^19, 62^. In addition, one study showed that AtREM1.3 from microsomal fractions is phosphorylated *in vitro* by AtCPK3 ^63^.

We therefore predicted that REM1.3 might share common functions with the evolutionarily conserved group 1b Arabidopsis REMs ^29^. AtREM1.2 and AtREM1.3 are close homologs to REM1.3 and group 1 *N. benthamiana* REMs (NbREMs) in term of protein sequence ^29, 31^ and they conserved at least the S74 and S91 phosphorylation sites ^34 35 48^ (Fig EV5A). Using super-resolution microscopy, Demir *et al.* showed that-when co-expressed in Arabidopsis leaves-REM1.3 and AtREM1.3 co-localized in the same PM-nanodomains ^64^. Importantly, transient expression of AtREM1.2 and AtREM1.3 in *N. benthamiana* epidermal cells impaired PVX-GFP cell-to-cell movement, as REM1.3 does (Fig EV5B), strengthening the hypothesis that the function of group 1 REMs might be conserved between homologs in different plant species ^31^.

We assayed the *in vitro* phosphorylation activity of the affinity-purified AtCPK3-GST towards the 6His-REM1.3, the 6His-REM1.3^N^ and the 6His-REM1.3^C^ as substrates. Similar to our previous results (Fig EV2B,C), AtCPK3-GST could phosphorylate strongly both 6His-REM1.3 and 6His-REM1.3^N^, but not 6His-REM1.3^C^ (Fig 5D). Addition of Ca^2+^ is essential for a strong kinase activity as shown by both kinase auto-phosphorylation and trans-phosphorylation (Fig 5D). AtCPK3-GST specifically phosphorylated S74 T86 and S91 residues of REM1.3, since the phosphorylation was abolished in the phosphomimetic mutant 6His-REM1.3^DDD^ (Fig 5E). These results suggest that AtCPK3 is a good candidate for group 1b REM phosphorylation and further support that the S74, T86, and S91 are the true phosphorylation sites of REM1.3 (Fig 3A, 5E).

### AtCPK3 interacts with group 1b REMs and restricts PVX propagation in a REM-dependent manner

CPKs harbor a variable N-terminal domain, a Ser/Thr kinase domain, an auto-inhibitory junction region and a regulatory calmodulin-like domain. The calmodulin-like domain contains four EF-hand binding motifs that determine the sensitivity of each kinase to calcium ^65 66^. To investigate the role of AtCPK3 in REM-dependent signaling, we generated AtCPK3 mutants presenting altered kinase activities. Deletion of the inhibitory junction region and the regulatory calmodulin-like domain in CPKs creates a constitutive active kinase while mutation of the aspartic acid residue in the catalytic center ‘DLK’ motif of the kinase domain to an alanine (D202A) creates a catalytically inactive or ‘dead’ kinase ^59^ (Fig 6A). We generated AtCPK3 full-length (AtCPK3), constitutive active (AtCPK3CA, 1-342) and kinase-dead (AtCPK3CA^D202A^) constructs for transient protein expression (Fig 6A). We evaluated their catalytic activities by expressing them transiently in Arabidopsis mesophyll protoplasts and performing immunoprecipitation coupled to kinase assays using 6His-REM1.3 and histone as a generic substrate ^59^. Autoradiography confirmed that *in vivo* purified AtCPK3CA-HA could trans-phosphorylate both 6His-REM1.3 and histone without the addition of calcium, while the point mutation D202A drastically abolished kinase activity (Fig EV6).

**Figure 6.**
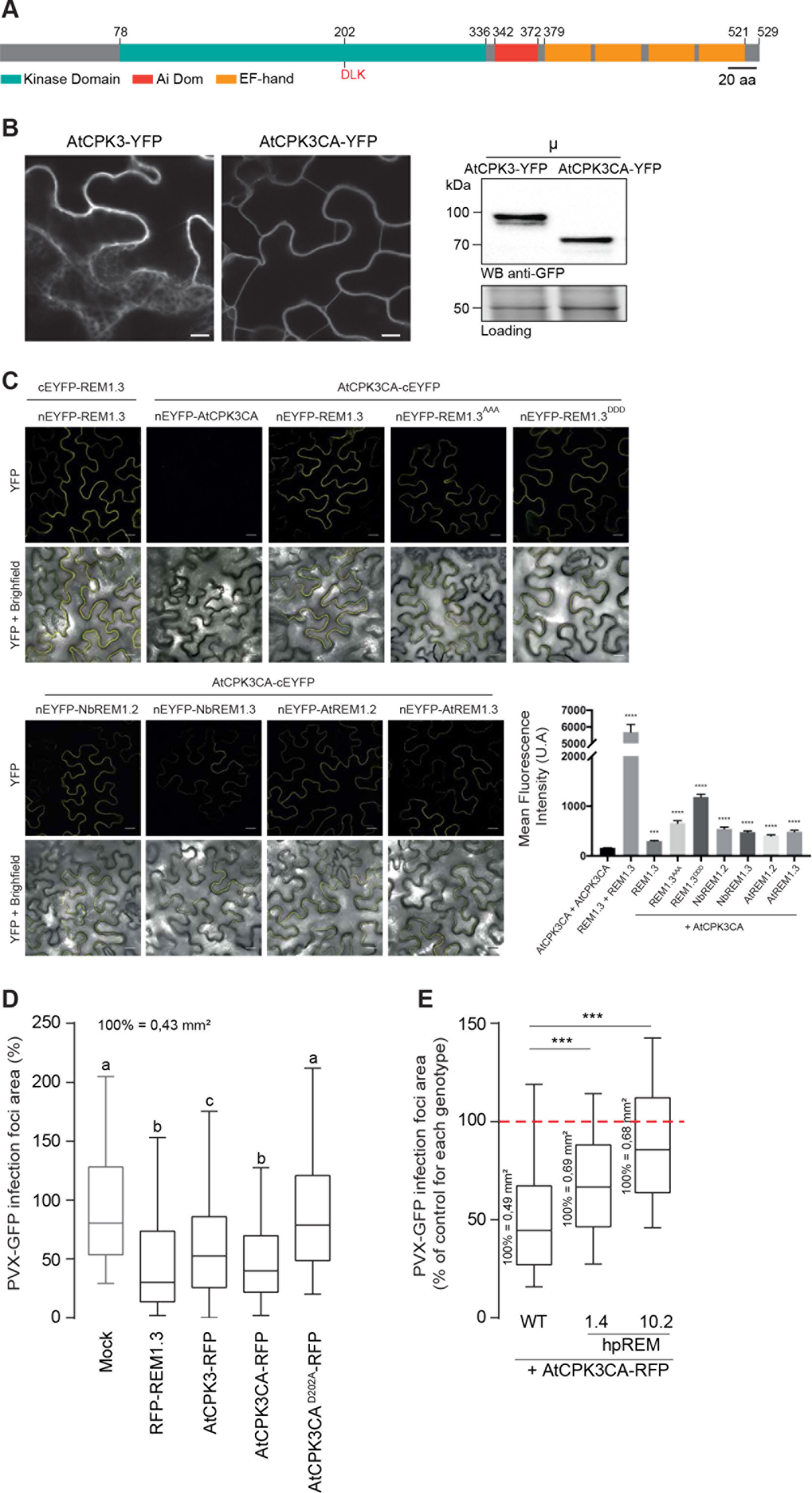
AtCPK3 physically interacts *in vivo* with group 1b REMs and impairs PVX cell-to-cell movement in a REM-dependent manner. (A) Primary sequence of AtCPK3. EF-hands are helix E-loop-helix F structural domains able to bind calcium. Ai : Autoinhibitory domain. The position of the DLK motif (Aspartic acid-Leucine-Lysine) at the catalytic domain conserved in all CPKs is indicated. (B) Confocal images showing AtCPK3-YFP and AtCPK3CA-YFP localization in *N. benthamiana* epidermal cells. Scale bar shows 10 μm. Western blot against GFP showing AtCPK3-YFP and AtCPK3CA-YFP expression in the microsomal fraction (μ) of *N. benthamiana* leaves. (C) *In planta* Bimo lecular Fluorescence Complementation (BiFC) analysis showing interaction of AtCPK3 with Group 1 REMs. REM1.3-YFP^N^/REM1.3-YFP^C^ was used as a positive control, and AtCPK3CA-nYFP/ AtCPK3CA-cYFP as a negative control. Mean fluorescence intensity at the cell boundary level was recorded using ImageJ. Statistical differences were determined by Mann-Whitney test compared to AtCPK3^CA^ +AtCPK3^CA^.*** p=0.0002, **** p ,0.0001. All scale bars indicates 20μm. (D) PVX-GFP spreading in *N. benthamiana* cells expressing RFP-REM1.3 or AtCPK3^FL^-RFP, AtCPK3^CA^-RFP, AtCPK3^CA D202A^-RFP Graph represents the area of PVX-GFP infection foci relative to the mock control (co-infiltration of PVX-GFP with empty *A. tumefaciens*). At least 200 PVX-GFP infection foci from at least 3 independent repeats were imaged at 5DAI. Letters indicate significant differences revealed by Dunn’s multiple comparisons test p<0.001․. (E) Effect of AtCPK3^CA^ on PVX-GFP cell-to-cell movement in WT *N. benthamiana* or in transgenic lines constitutively expressing hairpin REM (hpREM) constructs. At least 200 PVX-GFP infection foci from at least 3 independent repeats were imaged at 5DAI. For each *N. benthamiana* line the effect of AtCPK3^CA^ is expressed as a percentage of the corresponding mock control (empty Agrobacteria). Absolute values of the average foci area for each mock control are indicated.

We next examined the sub-cellular localization of both AtCPK3 and AtCPK3CA fused to YFP and found that both proteins disclosed a partial association with the PM, which was further confirmed by their presence, after cell fractionation, in the microsomal fraction at the expected molecular weight (Fig 6B) in good agreement with Mehlmer *et al.* ^63^. We further used AtCPK3CA to test the interaction with group 1b REMs. Bimolecular Fluorescence Complementation (BiFC) experiments showed that AtCPK3CA and REM1.3, REM1.3^AAA^ and REM1.3^DDD^ interact together at the level of the PM *in planta*. Importantly, we also confirmed the interaction of AtCPK3CA with homologous AtREM1.2 and AtREM1.3 (Fig 6C). REM1.3/REM1.3 interaction was used as a positive control, and AtCPK3CA /AtCPK3CA as a negative control.

We finally aimed to functionally characterize the AtCPK3- and REM1.3-mediated signaling in the context of PVX infection. Transient over-expression of AtCPK3-RFP alone induces a reduction of PVX-GFP infection foci, confirming that AtCPK3 is indeed important for antiviral responses in plant cells (Fig 6D). Expression of the constitutively-active AtCPK3CA-RFP had a stronger effect on PVX-GFP spreading and to a similar degree with the over-expression of REM1.3 alone (Fig 6D). AtCPK3’s function towards PVX movement was observed to be clearly mediated by its kinase activity, as the expression of the catalytically inactive mutant AtCPK3CA^D202A^ had no effect on PVX-GFP propagation (Fig 6D).

This raised the question whether the effect of AtCPK3CA on PVX propagation was REM-dependent. To tackle this question, we stably transformed *N. benthamiana* plants with a hairpin construct, to induce post-transcriptional gene silencing, which resulted in lowering RNA and protein expression of group 1 endogenous NbREMs (Fig EV7A,B). Consistent with previous studies ^7^, silencing of group 1 REM correlates with an increase of PVX cell-to-cell movement (Fig EV7C). Importantly, PVX assays demonstrated that AtCPK3CA ability to restrict PVX movement was impaired in two independent *N. benthamiana* lines underexpressing group 1 REM levels, namely lines 1.4 and 10.2 (with expression levels decreased respectively by 2 and 20 times) (Fig 6E), indicating that REMs might be the direct substrate of CPK3 *in vivo*. Altogether, these data suggest that CPK3 and group 1 REMs are major regulators involved in signaling and antiviral defense at the PM level.

## Discussion

Protein phosphorylation is a ubiquitous and specific mechanism of cell communication ^67^. The addition of a phosphate group on one or more critical residues of a given protein can induce important conformational changes that affect energetically favorable interactions and may lead to changes in its interacting network, localization, abundance and may influence the activity of protein signaling pools ^68^. Although, since the initial discovery of REM1.3 in 1989, accumulating evidence suggests that the functions of REM proteins are regulated by protein phosphorylation ^33–35^, the biological significance of this phosphorylation remained unclear to this date. REM proteins were among the first plant proteins described which supported the notion of PM subcompartmentalization to functional protein-lipid nanodomains ^7, 10 69^, also named membrane rafts ^2, 3^. In the present paper, we used REM1.3 and PVX as an experimental system to study the role of protein phosphorylation and membrane dynamics in the context of stress response.

### REM functions likely involve distinct PM compartments during plant PVX-sensing

Understanding how plants defend themselves against viruses remains a challenging field. The canonical plant immune response against viruses is mainly represented by the mechanism of RNA silencing ^70 71^, while additional mechanisms of plant antiviral defense involve hormonal signaling, protein degradation, suppression of protein synthesis and metabolic regulation ^42 70, 72^. Antiviral defense presents similarities to the immune response against microbes ^73 74 75^. Compelling evidence suggests that cell-surface as well as intracellular plant immune receptors recognize viral elicitors ^46 76 77 78 79 80 81^. An additional number of host cell components have been shown genetically to affect viral replication or cell-to-cell movement ^7 82^, indicating that more sophisticated plant defense mechanisms against viruses may exist.

For instance, manipulation of REM levels in transgenic *Solanaceae*, suggested that REM is as a positive regulator of defense against the PVX ^7, 13, 31^ by affecting viral cell-to-cell movement. We recently showed that REM1.3 does not interfere with the suppressor ability of PVX movement protein TGBp1, but specifically affects its gating ability ^30^. In this paper, we provide supporting mechanistic evidence that REM1.3 regulates the levels of callose accumulation at PD pit fields during PVX infection (Fig 1). Whether this function is mediated by a direct interaction with callose synthase/glucanase complexes remains however still unknown. Surprisingly, we found that REM1.3 is not dramatically recruited to PD pit fields, although its PD index is slightly increased after PVX infection (Fig 1). The spt-PALM VAEM microscopy data supports an increase of protein mobility and redistribution to distinct domains during PVX infection (Fig 1). These findings indicate the existence of a mechanism that operates at specific REM1.3-associated PM nanodomains, capable of regulating PD permeability (Fig 1). The dynamic partitioning between PM nanodomains and PD pit fields needs to be further studied.

### Plant PVX-sensing induces the activation of a calcium-dependent protein kinase

Since various studies have reported REM phosphorylation during plant-microbe interactions ^14 15 34 35^, we set out to address which kinase phosphorylates REM and whether REM1.3 phosphorylation plays a role in REM-mediated anti-viral defense. Indeed, our experimental findings show that plant PVX sensing induces the activation of a membrane-bound calcium-dependent protein kinase that in turn phosphorylates REM1.3 (Fig 2, Fig EV2, Fig 5). Importantly, we show that the kinase able to phosphorylate REM 1.3 is activated specifically by the expression of two PVX proteins, namely CP and TGBp1. Deciphering the exact mechanisms allowing the molecular recognition of those PVX components will be a crucial step toward understanding REM-mediated anti-viral defense.

Genetic studies have established that different CPKs comprise critical plant signaling hubs by sensing and translating pathogen-induced changes of calcium concentrations ^59 60^. Biochemical characterization of the kinase phosphorylating 6His-REM1.3 showed that its strong sensitivity to calcium (Figure 5C) corresponds to homologs of phylogenetic subgroup II CPKs from Arabidopsis ^59^. CPK3 is a prominent member of subgroup II, shown to function in stomatal ABA signaling ^83^, in salt stress response ^63 84^ and in a defense response against an herbivore ^85^. Interestingly, it was suggested that AtREM1.3 from taxonomical group 1 of REMs could be a substrate for AtCPK3 in response to salt stress ^63^. Here we show that AtCPK3 can interact *in vivo* with REM1.3 (Fig 6C) and that AtCPK3 phosphorylates REM1.3 in an *in vitro* kinase assay (Fig 5D). Transient overexpression of AtCPK3 in *N. benthamiana* resulted in a reduction of PVX propagation in a REM-dependent manner, providing compelling evidence that CPK3 together with REM contribute to the plant antiviral immunity. This is the first report demonstrating the participation of CPKs in plant basal immunity against viruses. The activation of CPKs by PVX supports the notion that PVX similarly to other pathogens or stress factors induces changes in calcium concentrations in the cell, which are sensed by the CPKs and translated via the phosphorylation of REM and other unknown downstream components. In *Nicotiana tabaccum* calmodulin isoforms are critical for the plant resistance against Tobacco Mosaic Virus and Cucumber Mosaic Virus, further illustrating the existence of virus-specific patterns of calcium signals ^86 87^. More work is needed to identify the CPK family members participating to the response and also the nature and specificity of those PVX-induced calcium changes.

### Phosphorylation regulates group 1 REM’s function during PVX cell-to-cell movement

AtCPK3 specifically phosphorylated REM1.3 at its N-terminal domain (residues 1-116), a domain displaying a mostly intrinsically disordered secondary structure (Fig 3A, 5). *In silico* analysis followed by a mutagenesis approach coupled with *in vitro* kinase assays revealed three major putative phosphorylation sites for REM1.3, namely S74, T86 and S91 on REM1.3. The *in vitro* phosphorylation of REM1.3 (Fig 3A, 5E) is almost totally lost when S74, T86 and S91 are mutated to non-phosphorylable residues, confirming these residues as major REM1.3 phosphorylation sites. The triple phospho-null mutant, YFP-REM1.3^AAA^, totally obliterated REM1.3’s capability to restrict PVX cell-to-cell movement. Reciprocally, REM1.3 triple phosphomimetic mutant YFP-REM1.3^DDD^ appeared fully functional (Fig 3D, 3E). These results strongly support the functional involvement of single or combined phosphorylation in the N-terminal domain of S74, T86 and S91 to establish REM’s function in the context of PVX infection. This is in contrast with LjSYMREM1 from *Lotus japonicus* which was shown to be phosphorylated at its C-terminal domain *in vitro* by SYMRK ^14^. Despite the fact that phosphorylation of REM proteins has been widely reported ^14 15 34 35 48^, this work firstly describes an associated role of REM-induced phosphorylation with its function.

### Toward the understanding of REMORIN function

Our finding that overexpression of *AtREM1.2* and *AtREM1.3* also restricts PVX-GFP cell-to-cell movement (Fig EV5B), suggests that REM phosphorylation and its associated functions might be conserved for some REMs of taxonomic group 1b. In good agreement, AtREM1.2 and AtREM1.3 localize to the same PM nanodomains ^64^ and maintain conserved phosphorylation sites with REM1.3 (Fig EV5A). By contrast, AtREM4.1 from subgroup 4 has an opposite effect against geminiviral propagation, promoting susceptibility to *Beet Curly Top Virus* and *Beet Severe Curly Top Virus* ^15^, but also does not present the same *in silico* phosphorylation profile ^48^ .

Overexpression of *REM1.3* restricts TMV propagation (Fig EV3A), and additionally hampers the gating activity of movement proteins from different virus genera ^30^, meaning that group 1 REM proteins might act as negative regulators for more viruses. These findings suggest that the initial hypothesis that REM1.3 causes the sequestration of the PVX virions at the PD ^7^ might not hold true, but rather that REM1.3 might have a more general role in plant stress and PD regulation (Fig 1,3). Interestingly, REM1.3 promotes susceptibility to *Phytophthora infestans* in *N. benthamiana ^37^*. The exact role of REM1.3 as a common regulator of different signaling pathways and its direct role in PD remain to be determined.

It has been speculated that phosphorylation in intrinsically disorder regions of proteins may act as a molecular switch and confer potential protein-protein interaction plasticity ^68, 88^. The intrinsically disordered REM1.3 N-terminal domain exhibits the most sequence variability in REM proteins, presumably conferring signaling specificity ^29, 48^. Phosphorylation of AtREM1.3’s N-terminal domain could stabilize coiled-coiled-associated protein trimerization and protein-protein interactions ^48^. Phosphorylated REM1.3 seems to be further targeted to PD-PM to trigger callose deposition. In good agreement, we found that the mobility in the PM of REM1.3 changed depending on its phospho-status (Fig 4). The triple phosphomimetic mutant exhibited a less confined and more mobile behavior at the PM, reminiscent of the WT protein in the context of PVX infection (Fig 4D). Similarly to the role of 14.3.3 proteins in plants ^89^, REM1.3 could act as a scaffolding protein, interacting with multiple members of a signaling pathway and tethering them into complexes to specific areas of the membrane. Hence, REM1.3 phosphorylation could act as a regulatory switch of protein conformations that would modulate REM1.3 specific interaction patterns and transient signalosomes at the PM. The triple phosphomimetic REM mutant might reflect a ‘functionally active’ form that constitutes REM-guided signalosomes against PVX-infection at the PM and should be exploited in future studies. The study of the phosphorylation-dependent interactions of REM1.3 (and related phosphocode) in regards to the modulation of REM1.3 PM dynamics and molecular function is the topic for future studies.

## Author contributions

AP performed the phosphorylation experiments. JG performed the spt-PALM experiments, PD index and callose measurements. JG and AP produced and imaged the REM mutants and built the figures. PG performed the BiFC assays. MB carried out the phosphorylation experiments with CPK3. AP, JG, AFD, VS performed the experiments of virus propagation. PG, AFD and VG produced the *N. benthamiana* hpREM-lines. MB, SGR, CZ, EEB and VG provided scientific expertise. AP, JG, SM and VG wrote the paper with the help of all the authors. VG and SM supervised the research. All authors read, edited, and approved the manuscript.

## Acknowledgments

Imaging was performed at the Bordeaux Imaging Center, member of the national infrastructure France BioImaging. AP was supported by the Greek fellowship program IKY in France. JG, PG and AFD are/were supported by the Ministère de l’Enseignement Supérieur et de la Recherche, France (MERS, doctoral grants). We acknowledge Dinesh Kumar for the gift of TMV-GFP clone, Alicia Zelada for the gift of PVXΔTGBp1 and viral protein constructs and Coralie Chesseron for greenhouse facilities. The IPS2 is supported by the LabEx Saclay Plant Sciences-SPS (ANR-10-LABX-0040-SPS). VG, SM, EEB, VS and VG are supported by the French ANR project “Potymove” (ANR-16-CE20-008-01). EEB, VG and SM are supported by the French ANR project “Connect”. CZ is supported by the Gatsby Charitable Foundation and the European Research Council (grant “PHOSPHinnATE”).

## Expanded View Figures (Supplementary figures)

**Fig EV1.**
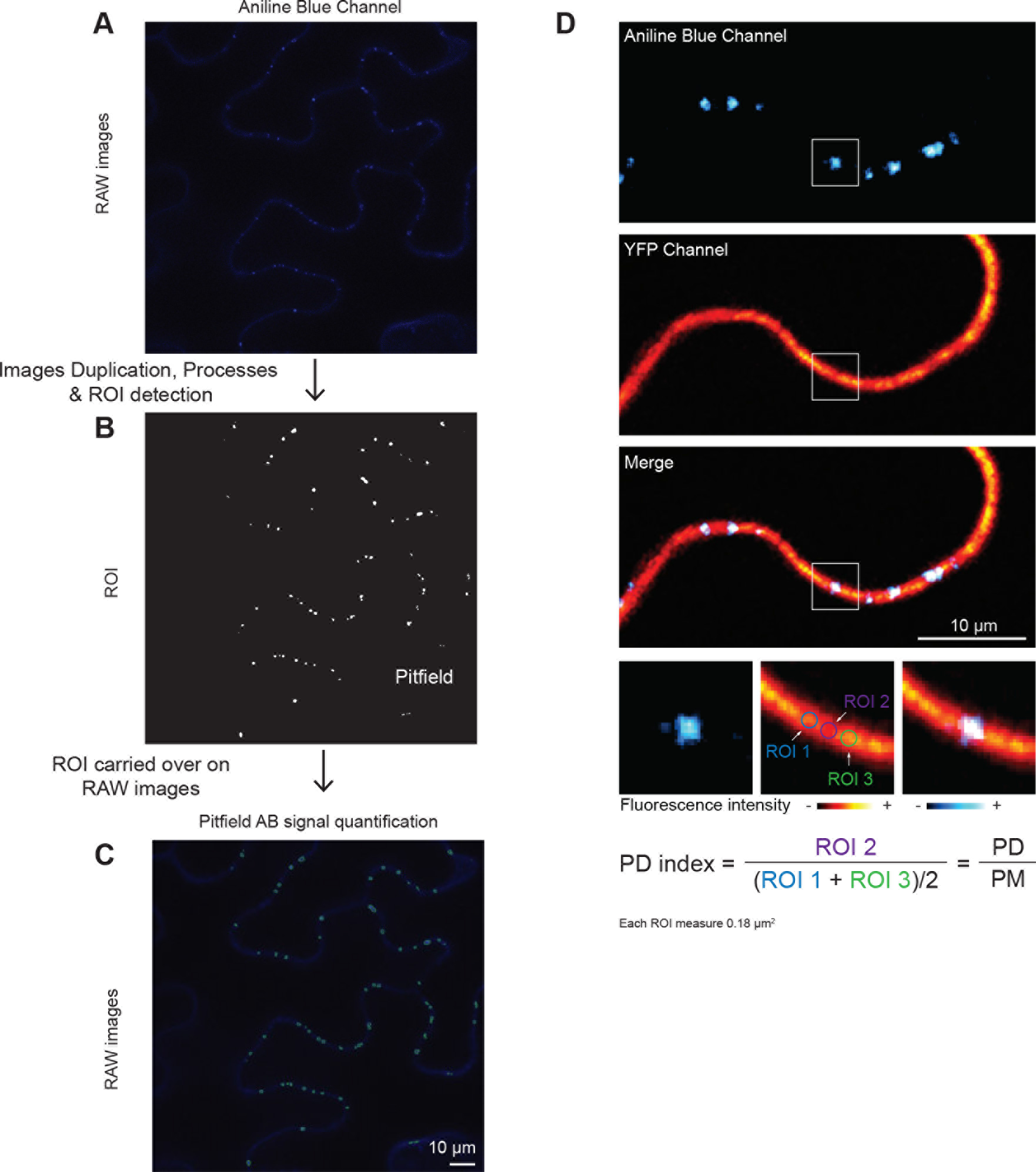
Callose quantification by aniline blue staining and PD index calculation. (A) Original sample image is an 8-bit, single-channel image. (B) Masks of total ROI objects before particle analysis were created using the following filters; background subtraction with a rolling ball radius as in ^90^; “smooth” twice and an auto-local threshold Max Entropy dark, creating a black and white mask, used for particle detection. (C) Overlay of outlines of the analyzed ROI (green; after particle analysis with particle size [3–100 pixel^2^ circularity [0.3–1] exclude on edge) with the original image. Scale bar indicates 10 μm. (D) Quantification of PD Index; after aniline blue labeled pit-field detection, YFP-REM1.3 fluorescence intensity was manually measured at pit-field level (ROI2) and surrounding PM (ROI1 and ROI3) using a circle of fixed area (0.18 μm^2^). The PD index was then calculated as the ratio between YFP-REM1.3 pit-field fluorescence (ROI2) and the mean of YFP-REM1.3 fluorescence intensity at surrounding PM (ROI1+ROI3).

**Fig EV2.**
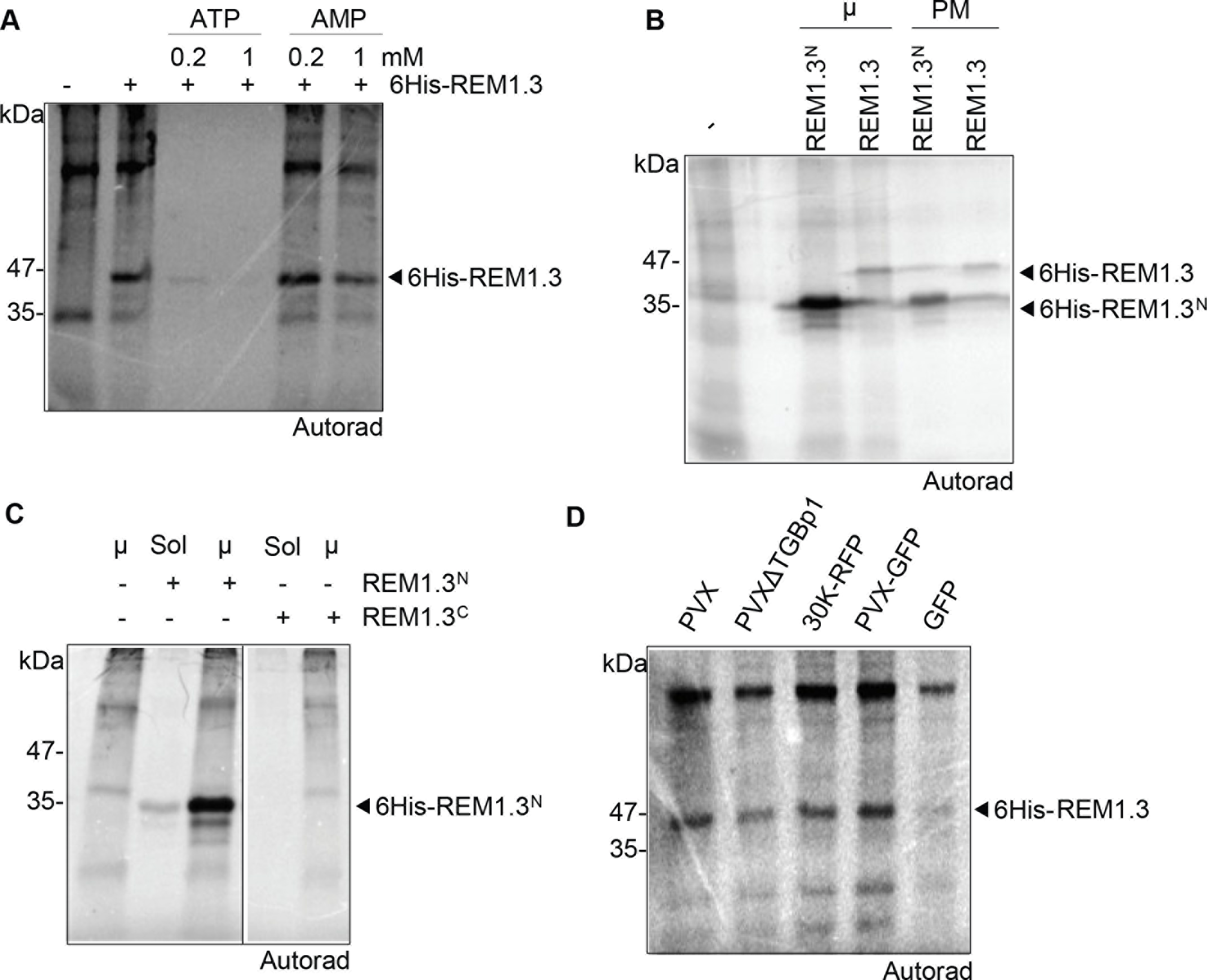
Analysis of *in vitro* 6His-REM1.3 phosphorylation. (A) Effect of the addition of ATP or AMP in *in vitro* phosphorylation assays of 6His-REM1.3 by kinase(s) in microsomal (μ) or PM extracts of *N. benthamiana* leaves developed by autoradiography. (B) 6His-REM1.3^N^ and 6His-REM1.3 phosphorylation by healthy *N. benthamiana* leaf microsomal (μ) and plasma membrane (PM) extracts. (C) 6His-REM1.3^N^ and 6His-REM1.3^C^ phosphorylation by kinase(s) in microsomal (μ) and soluble extracts. (D) 6His-REM1.3 was differentially phosphorylated by leaf microsomal extracts expressing the indicated constructs *i.e.* PVX alone, PVX deleted for TGBp1 (PVXPTGBp1), 30K protein from *Tobacco Mosaic Virus* (TMV), PVX fused to GFP, and GFP alone at 4 DAI. In all phosphorylation experiments about 10μg of total protein extracts and 1μg of affinity purified 6His-REM1.3, REM1.3^N^ or REM1.3^C^ were loaded per lane.

**Fig EV3.**
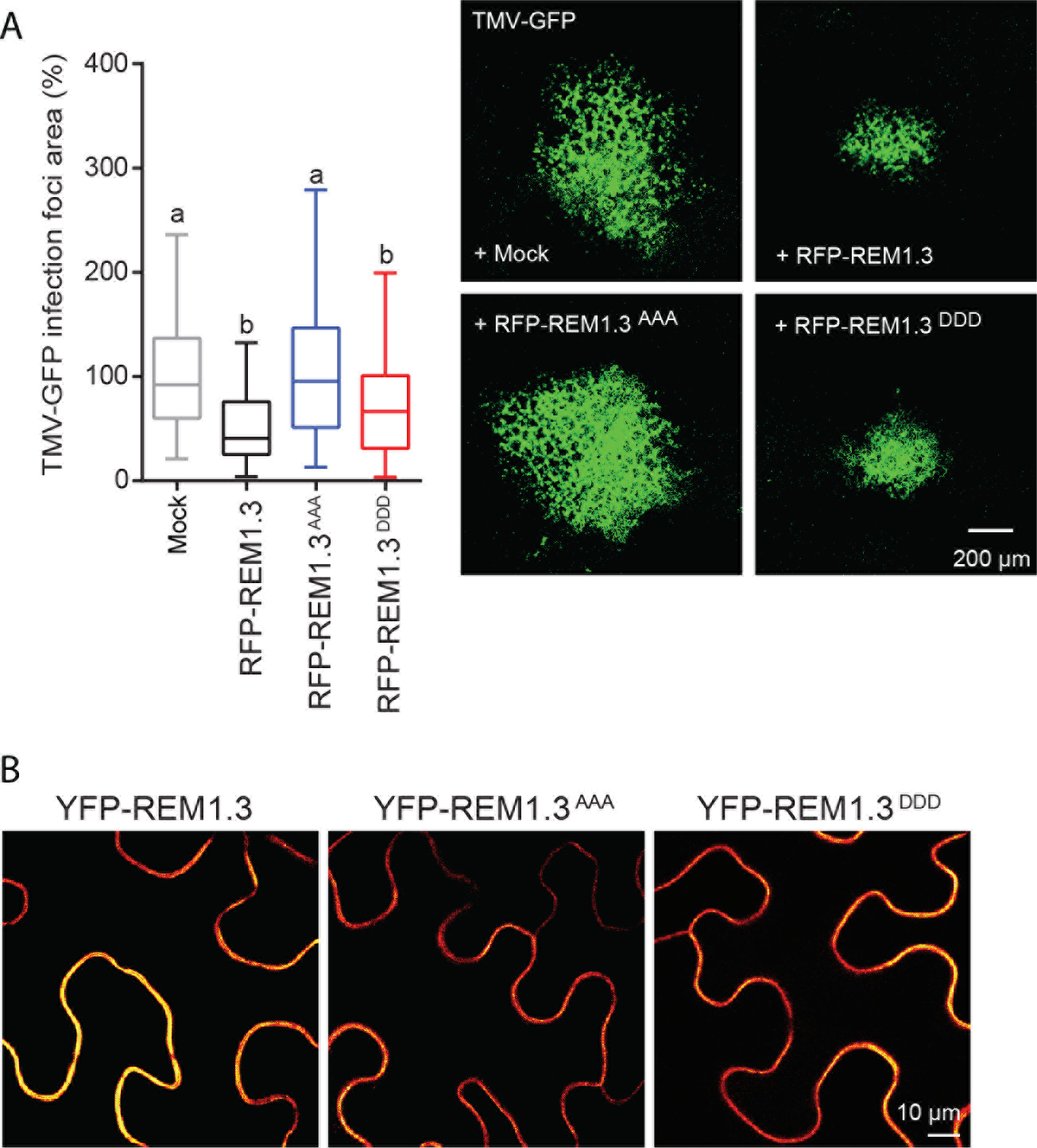
REM1.3 S74 T86 S91 phosphorylation is important to regulate Tobacco mosaic virus movement and modifies the protein lateral distribution in the PM. (A) Representative epifluorescence microscopy images of *Tobacco Mosaic Virus* (TMV-GFP) infection foci in *N. benthamiana* leaf epidermal cells at 5 DAI. Graph represents the relative foci area of REM1.3 or phosphomutants (S74, T86 and S91 into Alanine, AAA or Aspartic Acid, DDD) compared to mock control (co-infiltration of PVX-GFP with an empty *A. tumefaciens* strain). About 78–128 foci per condition were measured in 2 independent biological repeats. Dunn’s multiple comparison test was applied for statistical analysis, p<0.001. (B) Confocal microscopy images of secant views of *N. benthamiana* epidermal cells expressing YFP-REM1.3, YFP-REM1.3^AAA^ and YFP-REM1.3^DDD^ at 2 DAI. Scale bar indicates 10 μm.

**Fig EV4.**
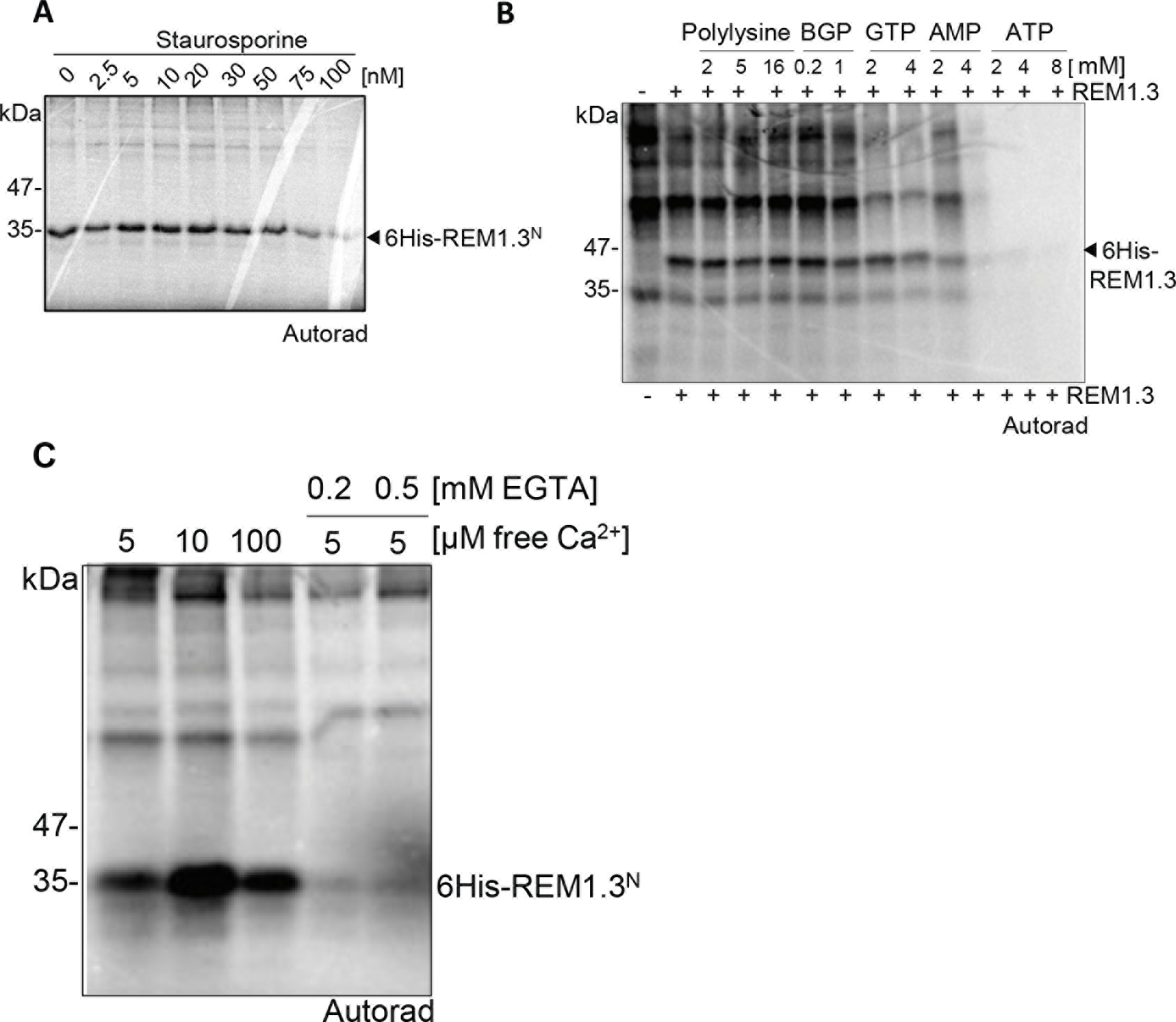
*In vitro* characterization of REM1.3 phosphorylation conditions. Autoradiography showing *in vitro* phosphorylation of purified 6His-REM1.3^N^ (A) or 6His-REM1.3 (B) by microsomal extracts of healthy *N. benthamiana* leaves in the presence of increasing concentrations of staurosporine (A) or Polylysine, β-glycerophosphate (BGP), GTP, AMP and ATP (B). (C) Effect of Ca^2+^ and EGTA on 6His-REM1.3^N^ phosphorylation by kinase(s) in microsomal extracts.

**Fig EV5.**
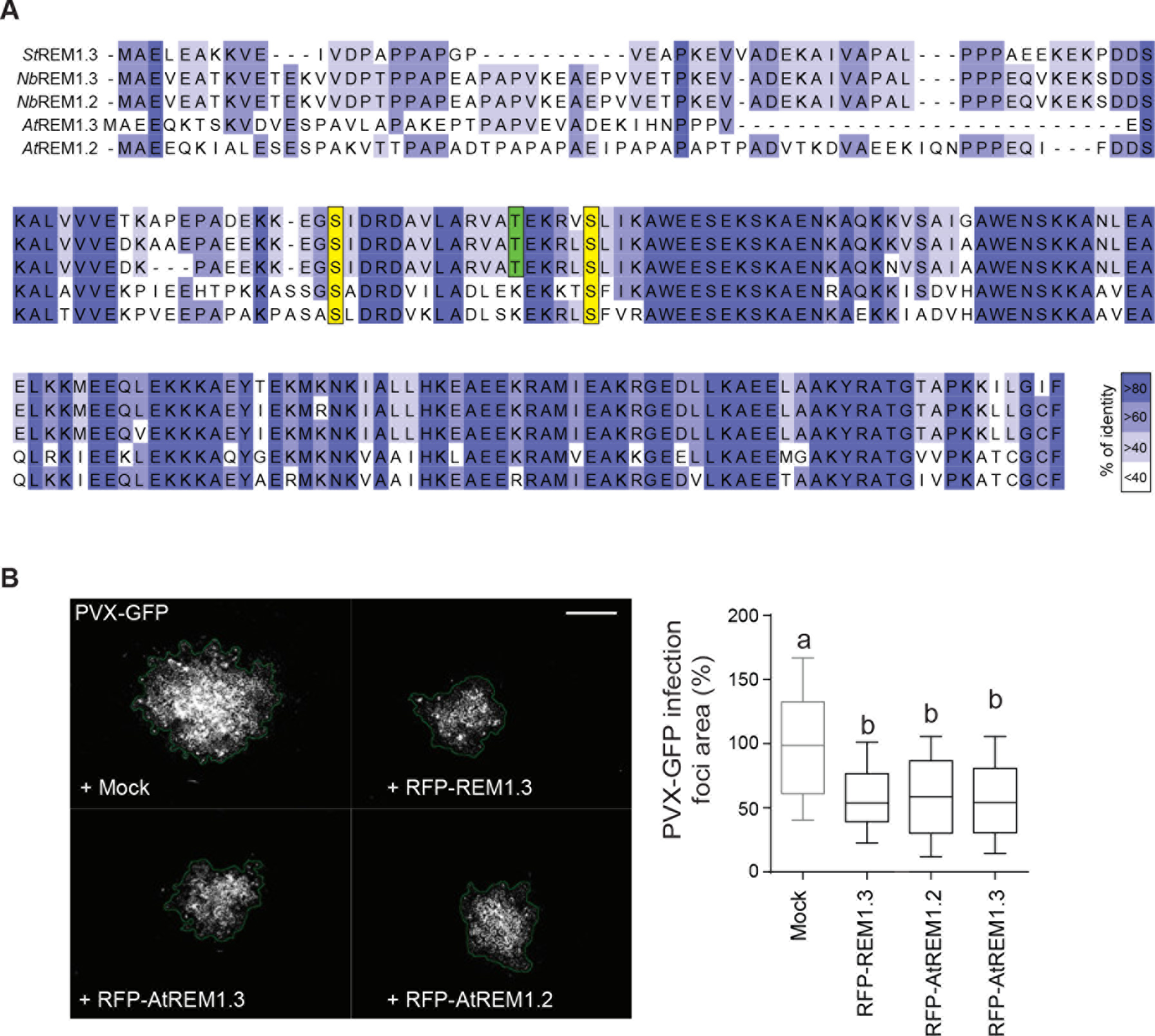
Group 1b AtREMs and REM1.3 have similar behavior against PVX cell-to-cell movement in *N. benthamiana* epidermal cells. (A) Clustal alignments of protein sequences of group 1b REMORINs: AtREM1.2, AtREM1.3, NbREM1.2, NbREM1.3 and REM1.3. Blue color-coding shows percentage of identity. The REM1.3 S74, T81 and S91 sites are highlighted. (B) *left*, Representative epifluorescence microscopy images of PVX-GFP infection foci on *N. benthamiana* leaf epidermal cells transiently expressing RFP-REM1.3, RFP-AtREM1.2 or RFP-AtREM1.3 at 5 DAI. Scale bar indicate 400 μm. *right*, Graph represents the relative PVX-GFP infection foci area in the presence of RFP-REM1.3 or Arabidopsis homologs compared to mock control (co-infiltration of PVX-GFP with empty *A. tumefaciens* strain). At least 184 foci per condition in 4 independent biological repeats were measured. Statistical differences are indicated by letters as revealed by Dunn’s multiple comparisons test pμ0.001.

**Fig EV6.**
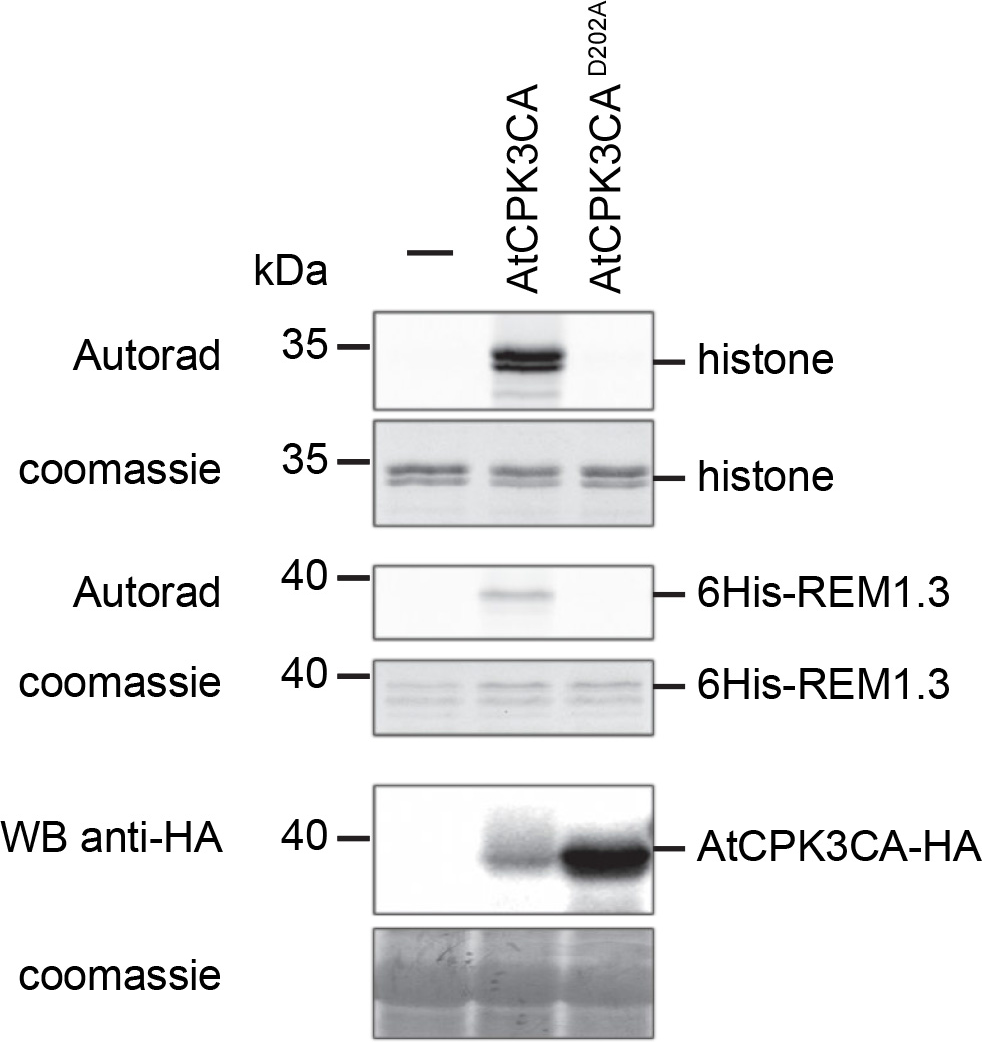
AtCPK3CA^D202A^ dead mutant does not phosphorylate REM1.3 *in* vitro. AtCPK3CA-HA and AtCPK3CA^D202A^-HA were expressed in *A. thaliana* mesophyll protoplasts. Immunoprecipitated proteins were incubated with ATP [γ-^33^P] and submitted to an *in vitro* kinase assay using 6His-REM1.3 or histone as substrates. *In vitro* kinase assays were revealed by autoradiography. Transphosphorylation of the substrates 6His-REM1.3 or histone is indicated. Western blot against HA shows the expression levels of the expressed proteins.

**Fig EV7.**
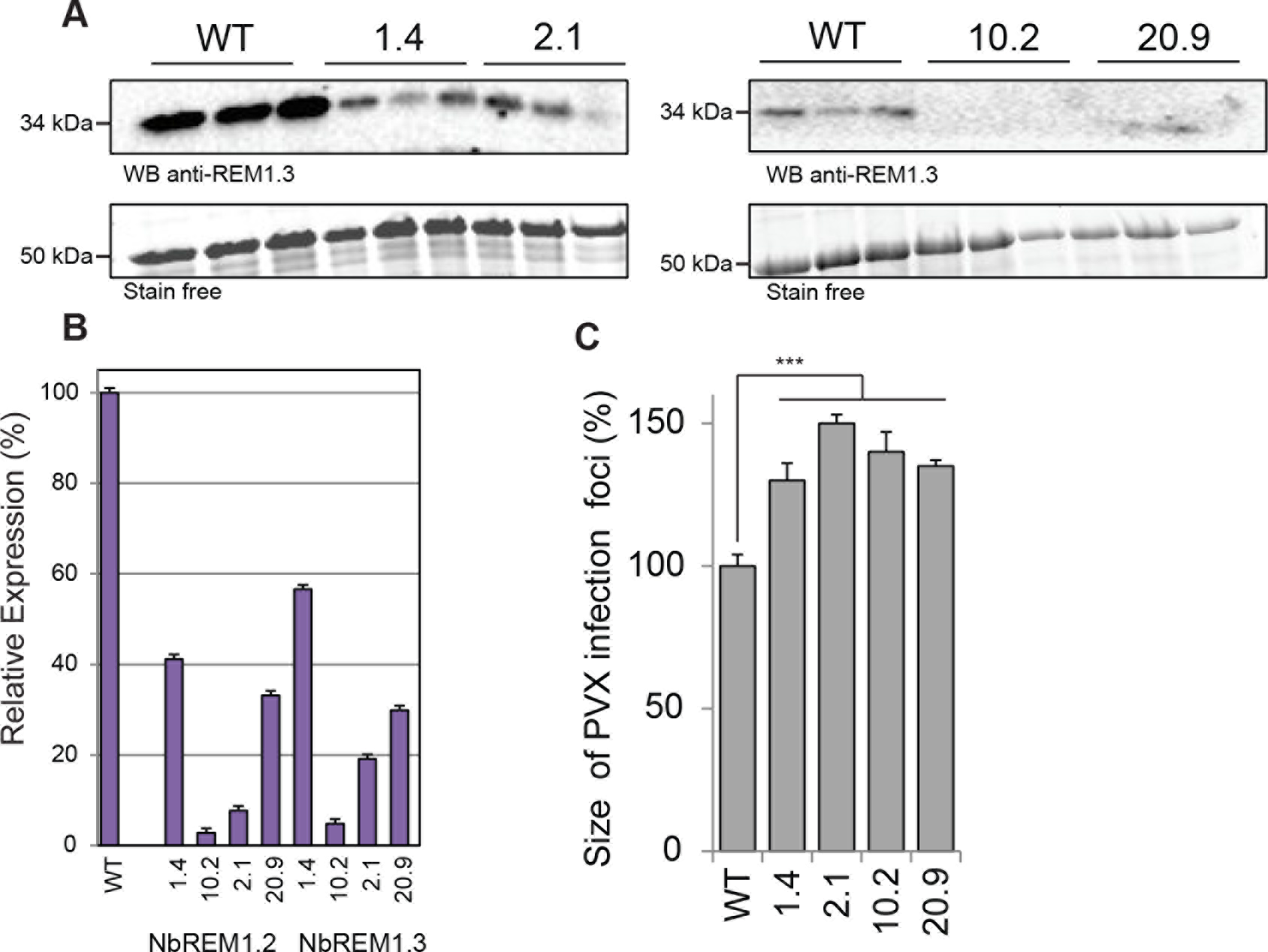
Stable transgenic lines *N. benthamiana* under-expressing group1 REMORINs. (A) Protein expression levels of endogenous NbREMs in the hpREM lines, determined by Western Blot analysis using anti-REM1.3 antibodies. Protein extracts from 3 independent plants per line were used. (B) Expression of endogenous NbREMs in the hpREM lines determined by RT-qPCR analysis. Results are expressed relative to the NbREMs expression levels in the WT background. RT-qPCR signals were normalized to actin levels. (C) PVX-GFP spreading is accelerated in the hpREM lines. Graph represents the PVX-GFP infection foci area in the different hpREM lines compared to WT. At least 3 independent experiments were performed. Error bars show +/− SEM. Statistical differences compared to WT were determined by Mann-Whitney test *** p<0.001.

## Materials and Methods

### Plant material

*Nicotiana benthamiana* plants were cultivated in controlled conditions (16 h photoperiod, 25 °C). Proteins were transiently expressed via *Agrobacterium tumefaciens*-mediated transformation for virus and PD functional assays as in ^13, 30^ or for the localization experiments as described in the appendix. For subcellular localization studies and protein extractions, plants were analyzed 2 DAI or at 4DAI in the phosphorylation assays. Imaging for PVX-GFP spreading assays and plasmodesmata GFP-diffusion experiments were done at 5 DAI. Details on molecular cloning and protein work, transgenic lines generation are described in the Appendix.

### Viral spreading and GFP diffusion assays

PVX-GFP cell-to-cell movement experiments were performed as previously described ^13, 31^, with minor modifications described in the Appendix. GFP diffusion at PD experiments was performed as previously described ^30^.

### Imaging

Live imaging and quantification were performed as described in ^31^. For Spt-PALM, images acquisitions and processing were done as previously described ^91^. PD index was calculated as in ^25^. More details described in the Appendix.

### *In vitro* kinase assays

For the *in vitro* REM1.3 phosphorylation assays about 2 μg of total plant extracts were incubated with 1 μg of affinity-purified 6His:REM1.3 protein variants for 10 minutes at room temperature and in a phosphorylation buffer (Tris-HCl 30mM, EDTA 5mM, MgCl_2_ 15mM, DTT 1mM, Na_3_VO_4_ 2,5 mM, NaF 10 mM and 10 μCi/reaction ATP [γ-^33^P]- (3000Ci/mmol, Perkinelmer)). The buffer contained also 10^-5^ M or gradual concentrations of free Ca^2+^ as in ^92^. Reactions were performed for 15 minutes in a volume of 25 μl. The reactions were stopped by the addition of 15μl of 6x Laemmli buffer. Proteins were separated by SDS/PAGE and phosphorylation status of REM1.3 was analysed by autoradiography using a phosphor-Imager and quantified by ImageQuant TL program. AtCPK3 variants phosphorylation assays were performed as described in ^93^.

## Appendix Supplementary Methods

### Cloning and molecular constructs

All vectors constructs were generated using classical Gateway^®^ cloning strategies (http://www.lifetechnologies.com), pDONR211 and pDONR207 as entry vectors, and pK7WGY2, pK7YWG2, pK7WGR2, pK7RWG2, and pGWB14 and pGWB15 as destination vectors (Miyazawa *et al.* 2010). The REM1.3^1-116^, REM1.3^117-198^ and REM1.3^S74A T86A S91A^, REM1.3^S74D T86D S91D^ were synthesized in a pUC57 vector (including the AttB sites) by Genscript (http://www.genscript.com/) and next cloned to Gateway^®^ destination vectors. AtCPK3^D202A^ mutant was generated by site-directed mutagenesis as previously described ^94^ with minor modifications. For BiFC experiments, the genes of interest were cloned into pSITE-BIFC- C1nec, -C1cec, -N1nen, and -N1cen destination vectors ^95^.

### Generation of transgenic stable hairpin REM *N. benthamiana* lines

Leaf discs were cut from *N. benthamiana* leaves, transferred on petri plates containing culture medium (complete Murashige and Skoog medium (MS) supplemented with 30g/L saccharose, pH 5,8; phytoagar HP696 (Kalys) 5,5 g/L and the hormones: AIA 0,1 mg/L, BAP 2 mg/L) and incubated for 48 h in the growth room (16 h photoperiod, 30 μmol photons.m2.s^−1^, 23 °C). For the transformation, the *N. benthamiana* plants disk leaves were incubated with the *Agrobacterium* cultures (GV3101 strain) carrying the plasmid of interest for 20 min. The leaf samples were next placed on plates with the complete medium previously described. 48 hours later, the leaf fragments were washed 3 times with sterile water (with 0,1 $ Tween_20_). The leaf fragments were next washed with MS complete medium supplemented with Timentin (300 μg/mL). The leaves were next placed on plates with regeneration medium (MS culture medium, as previously described, supplemented with 300 mg/L of timentin and 150 mg/l of kanamycin). The plates were next incubated in the growth room. The explants were transferred to fresh regeneration medium with a maximum periodicity of 7 days until the development of callus. The regenerated seedlings were transferred to a rooting medium (MS, sucrose 30 g/L, phytoagar 5,5 g/L, timentin 200 mg/L, kanamycin 150 mg/L). The regenerated plants (T0) were transferred to the greenhouse for growth and self-fertilization. Homozygous T2 lines carrying a single transgene insertion were selected by segregation analysis on selective Kanamycin media and used for physiological studies and phenotypic characterization. The expression or silencing levels of the YFP-REM1.3 or hpREM lines respectively was controlled by cytological, biochemical and expression analysis.

### Viral spreading, GFP diffusion assays

To assess spreading of PVX-GFP in tobacco leaves, *A. tumefaciens* strain GV3101 carrying the constructs tested were infiltrated at a final optical density at 600 nm (OD_600nm_) = 0.2 together with the same strain carrying the plasmid pGr208, which expresses the PVX:GFP complementary DNA harboring GFP placed under the control of a Coat protein promoter, as well as the helper plasmid pSoup ^96^ at final OD_600nm_ of 0.001. Viral spreading of PVX-GFP was visualized by epifluorescence microscopy (using GFP long pass filter on a Nikon Eclipse E800 with x4 objective coupled to a Coolsnap HQ2 camera) at 5 DAI and the area of at least 30 of PVX-GFP infection foci was measured using Fiji software (http://www.fiji.sc/) via a homemade macro or ImageJ. The expression levels of transiently expressed constructs were confirmed by Western blot. All the experiments were repeated at least three times.

### Transient expression in *N. benthamiana*

Four-week-old *N. benthamiana* greenhouse plants grown at 22-24 °C were used for *Agrobacterium tumefaciens*-mediated transient expression. *A. tumefaciens* were pre-cultured at 28 °C overnight, and used as inoculum for culture at initial OD_600nm_ of 0.15 in pre-warmed media. Cultures were grown until OD_600nm_ reached 0.6 to 0.8 values (3-5 h). Cultures were then centrifuged at 3,200 g for 5 min, pellet were washed twice using water to the desired OD_600nm_. Bacterial suspensions at OD_600nm_ of 0.2 and 0.1 were used for subcellular localization and Spt-PALM experiments, respectively. The bacterial suspensions were inoculated using a 1-mL syringe without a needle by gentle pressure through a ,1mm-hole punched on the lower epidermal surface ^97^. Transformed plants were incubated under normal growth conditions for 2 days at 22-24 °C. Transformed *N. benthamiana* leaves were analyzed 2-5 DAI depending on the experiment.

### Epidermal cells live imaging and quantification. Bimolecular Fluorescence Complementation

Live imaging was performed using a Leica SP5 confocal laser scanning microscopy system (Leica, Wetzlar, Germany) equipped with Argon, DPSS and He-Ne lasers and hybrid detectors. *N. benthamiana* leaf samples were gently transferred between a glass slide and a cover slip in a drop of water. YFP and mCitrine (cYFP) fluorescence were observed with similar settings (*i.e.*, excitation wavelengths of 488 nm and emission wavelengths of 490 to 550 nm). In order to obtain quantitative data, experiments were performed using strictly identical confocal acquisition parameters (*e.g.* laser power, gain, zoom factor, resolution, and emission wavelengths reception), with detector settings optimized for low background and no pixel saturation. Pseudo-colored images were obtained using the “Red hot” look-up-table (LUT) of Fiji software ( http://www.fiji.sc/). All quantifications were performed for at least 10 cells, at least two plants by condition with at least 3 independent replicates. BiFC images were taken 2DAI by confocal microscopy (Zeiss LSM 880). Quantification of fluorescent intensities was performed by ImageJ.

### Spt-PALM, single molecule localization and tracking

*N. benthamiana* epidermal cells were imaged at room temperature (RT). Samples of leaves of 2-week-old plants expressing EOS constructs were mounted between a glass slide and a cover slip in a drop of water to avoid dehydration. Acquisitions were done on an inverted motorized microscope Nikon Ti Eclipse (Nikon France S.A.S., Champigny-sur-Marne, France) equipped with a 100× oil-immersion PL-APO objective (NA = 1.49), a TIRF arm, a Perfect Focus System (PFS), allowing long acquisition in oblique illumination mode, and a sensitive Evolve EMCCD camera (Photometrics, Tucson, USA). Images acquisitions and processing were done as previously described ^91^.

### *In silico* analysis of REM1.3 protein sequence

Prediction of putative phosphorylation sites was performed by Diphos, DEPP and NETPHOS coupled with published data. Disordered domains were performed by pDONR VL XT.

### *In vitro* CPK3 kinase assays

CPK3-HA was transiently expressed in mesophyll protoplasts and immunopurified with anti-HA antibodies as performed in ^93^ while CPK3-GST recombinant protein was purified from bacterial extracts as reported in ^61^. For *in vitro* kinase assays, the tagged CPK was incubated with 0.5-1 μg histone or 6His-REM1.3 proteins in the following kinase reaction buffer (20 mM Tris HCl pH 7.5, 10 mM MgCl_2_, 1 mM DTT, 50 μM cold ATP, ATP [γ-^33^P] 2 μCi per reaction, 1 mM CaCl_2_ or 5 mM EGTA) in a volume of 15 μL for 30 min at RT. The reaction was stopped with 5 μL 4X Laemmli buffer, then samples were heated at 95 °C for 3 min. Proteins samples were separated by SDS-PAGE on 12% acrylamide gel. After migration, the gel was dried before exposing against a phosphorScreen to reveal radioactivity on a Storm Imaging system (GE Heathcare). The gel was then rehydrated for Coomassie staining.

### Protein Work

SDS/PAGE and Western Blot analysis, protein extractions and recombinant protein purification were performed as in ^13^. Cell fractionation and extractions followed the established protocol from ^50^ and ^53^.

### Accession Numbers

Sequence data from this article can be found in the Arabidopsis Genome Initiative (https://www.arabidopsis.org/index.jsp), and GenBank/EMBL (https://www.ncbi.nlm.nih.gov/genbank/) databases under the following accession numbers: REM1.3 (NP_001274989), AtREM1.2 (At3g61260), AtREM1.3 (At2g45820), AtCPK3 (At4g23650).

